# Fear Conditioning Potentiates the Hippocampal CA1 Commissural Pathway In vivo and Increases Awake Phase Sleep

**DOI:** 10.1101/2020.05.13.093237

**Authors:** Manivannan Subramaniyan, Sumithrra Manivannan, Vikas Chelur, Theodoros Tsetsenis, Evan Jiang, John A. Dani

## Abstract

The hippocampus is essential for spatial learning and memory. To assess learning we used contextual fear conditioning (cFC), where animals learn to associate a place with aversive events like foot-shocks. Candidate memory mechanisms for cFC are long-term potentiation and long-term depression, but there is little direct evidence of them operating in the hippocampus in vivo following cFC. Also, little is known about the behavioral state changes induced by cFC. To address these issues, we recorded local field potentials in freely behaving mice by stimulating in the left dorsal CA1 region and recording in the right dorsal CA1 region. Synaptic strength in the commissural pathway was monitored by measuring field excitatory postsynaptic potentials (fEPSPs) before and after cFC. After cFC, the commissural pathway’s synaptic strength was potentiated. Although recordings occurred during the wake phase of the light/dark cycle, the mice slept more in the post-conditioning period than in the pre-conditioning period. Relative to awake periods, in non-rapid eye movement (NREM) sleep the fEPSPs were larger in both pre- and post-conditioning periods. We also found a significant negative correlation between the animal’s speed and fEPSP size. Therefore, to avoid confounds in the fEFSP potentiation estimates, we controlled for speed-related and sleep-related fEPSP changes and still found that cFC induced long-term potentiation, but no significant long-term depression. Synaptic strength changes were not found in the control group that simply explored the fear-conditioning chamber, indicating that exploration of the novel place did not produce the measurable effects caused by cFC. These results show that following cFC, the CA1 commissural pathway is potentiated, likely contributing to the functional integration of the left and right hippocampi in fear memory consolidation. In addition, the cFC paradigm produces significant changes in an animal’s behavioral state, which are observable as proximal changes in sleep patterns.

## Introduction

Learning and memory are vital for the survival of animals, and therefore disorders affecting these systems are debilitating. Contextual fear conditioning (cFC) is an associative learning paradigm and a commonly used model of clinical anxiety disorders (Maren et al., 2013; Shin & Liberzon, 2010). In this task, animals learn within a single 5-10 minute trial to associate a distinct place with foot-shocks. Thus, this assay enables the observation of neural circuit changes across distinct learning phases (e.g. memory acquisition, consolidation, and recall) on a relatively brief timescale. Learning in this task requires the hippocampus (Maren et al., 2013), whose well-defined synaptic pathways are amenable to experimental manipulations. By leveraging the tractability of hippocampal pathways and the cFC behavioral assay, we examined real-time neural circuit changes associated with fear memories. Ideally those pathways may provide targets for the ongoing development of treatments aimed at relieving stress-related memory disorders.

The mechanisms of associative fear memory are complex and involve molecular to systems-level changes in the brain, and fully understanding these mechanisms will require integrating information from various levels of neural function (Sporns, 2011). Most studies to date have focused on either the systems-level or the molecular/structural level. At the systems-level, recent studies have focused on associative fear memory-induced changes in inter-regional coupling (Girardeau et al., 2017; Narayanan et al., 2007; Popa et al., 2010; Seidenbecher et al., 2003), neuronal oscillations and synchrony (Lesting et al., 2013; Likhtik et al., 2013; Ognjanovski et al., 2014; Ognjanovski et al., 2017; Pape et al., 2005), engram (Tonegawa et al., 2015) and place field formation (Moita et al., 2004; Wang et al., 2012). At the cellular, molecular and structural level, studies have focused on changes in intracellular signaling, receptor and ion channel function and neuronal morphology that are associated with fear memory formation (Buffington et al., 2014; Johansen et al., 2011; Karunakaran et al., 2016; Nicoll, 2017; Penn et al., 2017). These studies imply that these molecular/structural changes should lead to systems-level changes, and the link between these levels should be circuit-level changes, which manifest as changes in synaptic strength. While a handful of seminal studies have shown that learning-related changes in synaptic strength do occur in vivo in some aversive learning paradigms (Broussard et al., 2016; Gruart et al., 2006; Whitlock et al., 2006), this crucial link has not been established for cFC.

The hippocampal commissural pathways are highly conserved during evolution (Suárez et al., 2014), and they strongly interconnect the subregions of the left and right hippocampi (Laurberg, 1979; Swanson et al., 1978). However, the functions of these connections are only beginning to be unraveled (Phelps et al., 1991; Schimanski et al., 2002; Zhou et al., 2017). Although in vivo LTP can be artificially induced in these pathways (Bliss et al., 1983; Martin et al., 2019; Stäubli & Scafidi, 1999), it is unknown if plasticity occurs in vivo in these pathways in association with cFC. Moreover, unlike artificially induced plasticity, learning-related changes are widely distributed (Eichenbaum, 2016), making their detection harder at the circuit level. Hence, although fear conditioning-related plasticity likely occurs in the commissural pathways, it is unknown if it will be detectable in vivo. Also unknown is the synaptic plasticity’s temporal profile, which may be very different from that resulting from artificial plasticity induction owing to potential changes in the animal’s behavioral states accompanying learning. To characterize how the commissural pathway’s strength changes in association with behavioral learning, we periodically monitored the synaptic strength of the dorsal CA1 commissural pathway before and immediately after the cFC task. One impediment to assessing learning-related synaptic changes in this pathway is that behavioral states such as wakeful voluntary motion, still alertness, REM sleep, and NREM or slow wave sleep, are also known to influence excitability and synaptic transmission in the hippocampus (Buzsaki et al., 1981; Green et al., 1990; Grosmark et al., 2012; Hulse et al., 2017; Segal, 1978; Winson & Abzug, 1977, 1978a, 1978b). In addition, the speed of an animal’s movement is also known to be correlated with fEPSP response size (Kemere et al., 2013). To isolate learning-related effects from speed-related and potential behavioral state-related effects on synaptic plasticity measurements, we monitored the animal’s movement and the gross behavioral states using motion tracking and power spectral analysis of hippocampal recordings.

Our analyses showed that following cFC, behavioral states changed: mice slept more after the cFC although they were in the wake phase of their light/dark cycle. As reported previously in rats (Kemere et al., 2013), we found a significant negative correlation between animal speed and fEPSP size in mice. During the NREM part of these sleep periods, the fEPSPs were larger in amplitude. We accounted for the speed- and NREM-related potential confounds when measuring the cFC-induced synaptic changes. Although we did not find evidence for long-term depression, our results showed that the dorsal CA1 commissural pathway undergoes significant in vivo long-term potentiation in real-time following cFC.

## Results

To determine whether long-lasting synaptic changes occur in the hippocampal synapses due to associative learning, we measured synaptic strength in the commissural pathway of the dorsal CA1 region in freely moving mice (Figure 1A). We placed a stimulating electrode in the left CA1 and positioned one or more recording electrodes in the right CA1 (Figure 1B). With this configuration, the stimulation response in the right CA1 arises from monosynaptic activation of direct commissural projections from the left CA1 (Zhou et al., 2017) and also likely from a disynaptic route where action potentials from the left CA1 antidromically activate the left and right CA3 which then activates the right CA1. Synaptic strength was quantified by periodically measuring the slope of the population field EPSP (fEPSP) arising from brief electrical stimulation (Figure 1C-D). All experiments were done during the awake (dark) phase of the animals’ light/dark cycle (Figure 1E). During recordings, mice remained undisturbed in their home cage that was placed inside a sound-attenuating recording chamber. After a 2h baseline measurement, mice (n = 24) were removed from their home cage and subjected to cFC (Figure 1E). To examine the conditioning-related neural changes immediately following fear conditioning, synaptic strength was measured again for another 3h. At least one day after these recordings, fear memory was tested by leaving the animals in the original conditioning chamber for 5 min and measuring the percentage of time spent in freezing. We did this test 24h post-conditioning in one group of mice (n = 12) and 1-week post-conditioning in a second group of mice (n = 12). Both groups showed significant memory retention (i.e., freezing behavior) suggesting that the conditioning protocol effectively induced long-lasting memory: p < 10^-5^ for both groups, t-test; % freezing after 24h, 37 ± 4%; after 1-week, 49 ± 5%; mean ± 1SE. There was no significant difference between the two groups (p = 0.058, t-test). The pooled group showed a significant fear memory (Figure 1F): % time spent in freezing during the first 2 min pre-shock exploration period on conditioning day was 0.2 ± 0.1%; % freezing on testing day was 43 ± 3%, n = 24, p < 10^-5^, paired t-test on conditioning day versus testing day freezing level.

**Figure 1.**
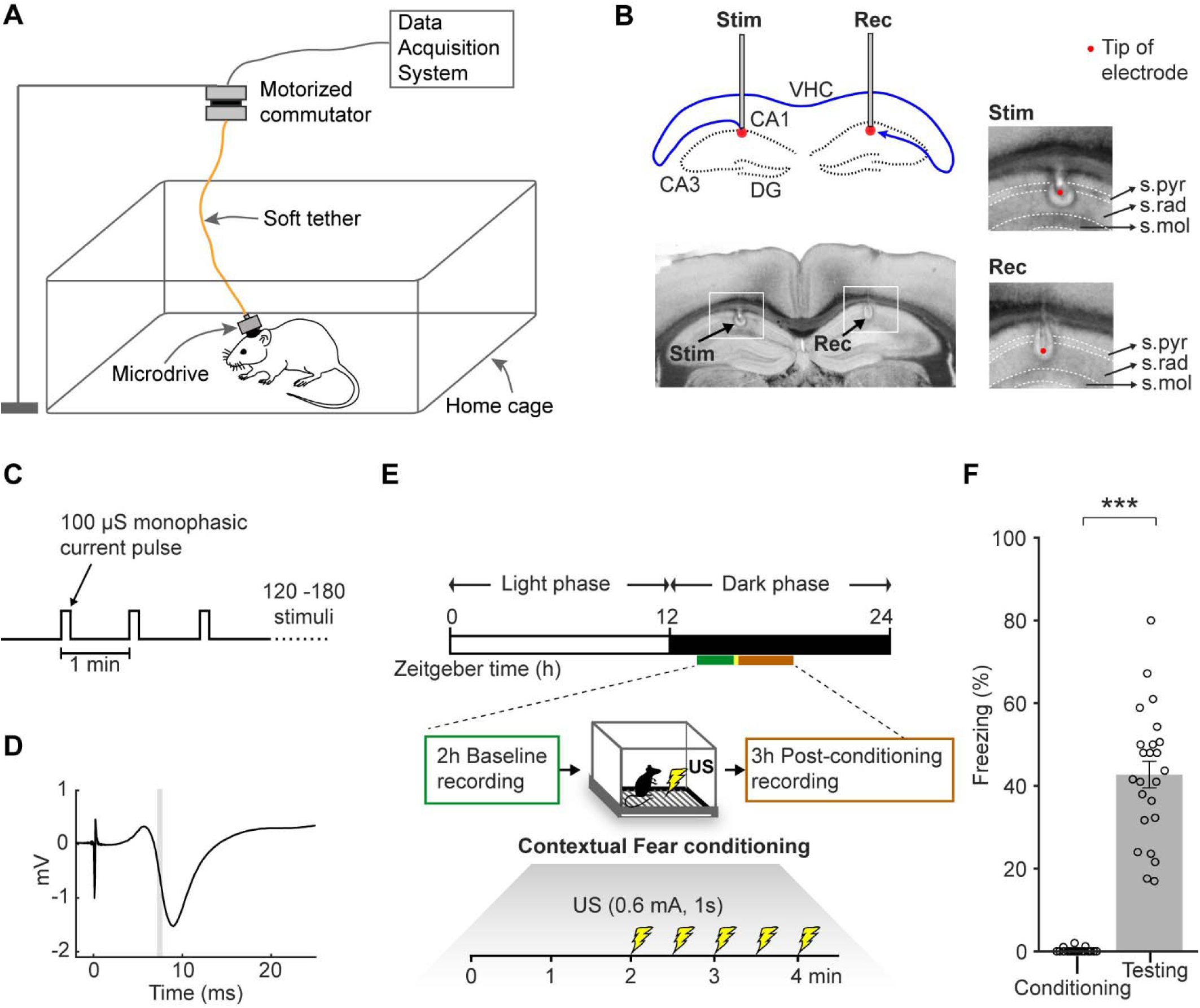
The contextual fear conditioning (cFC) setup for measuring the in vivo synaptic strength changes in the dorsal CA1 commissural pathway. ***A***, Illustration of neural recordings with a microdrive carrying a stimulating electrode and multiple recording electrodes. The contents of the home cage, such as food, bedding and nest, are not depicted. ***B***, Top left: Schematic drawing showing the location of the stimulating (stim) and the recording (rec) electrodes, and the CA1-to-CA1 commissural pathway (blue) coursing through the ventral hippocampal commissure (VHC). Cell layers of CA1, CA3, and dentate gyrus (DG) are drawn as black dotted traces. Bottom left: Coronal brain section showing the stimulating and a recording electrode location in the CA1 identified by the electrolytic lesions (“halo” around the electrode tips marked by black arrows). Insets on right show expanded views of the stimulating and recording electrode locations with a red dot indicating the tip. Dotted white lines set the boundaries of *stratum pyramidale* (s.pyr), *radiatum* (s.rad), and *moleculare* (s.mol). ***C***, Stimulation protocol applied to the left CA1, which was stimulated every 1 min with 100 μs square pulses for 2-3 h. ***D***, Field EPSP response from the recording electrode shown in ***B***. Time zero marks the onset of the stimulus pulse. The slope measured in a 0.5 ms window (gray box) of the downward deflecting portion of the response was taken as a measure of synaptic strength. ***E***, Illustration of the cFC procedure. Recording timelines are represented on the Zeitgeber time scale (Zeitgeber time represents time (h) from the onset of house light). Experiments were conducted during the awake, dark phase. In the home cage, the synaptic strength was periodically measured as in ***C*** for 2 hours before (baseline), and 3 hours immediately after cFC. The green and orange bars under the Zeitgeber time scale represent baseline and post-conditioning recording periods respectively. The baseline measurement started ∼1.5 h after the house light was turned off. US: unconditioned stimulus. ***F***, Percentage of time spent in freezing, measured on conditioning day (Conditioning) and on testing day (Testing). Error bars show SEM, and *** p < 10^-5^ is the significant difference between conditioning and testing day freezing levels.

Stimulation and recording sites were restricted to the dorsal CA1 region (Figure 2A). We consider two scenarios on how the learning-related changes might be distributed across the region sampled by these recording electrodes. The synaptic changes could be distributed evenly throughout a large region of the hippocampus. In this case, all the electrodes will report either a net potentiation, depression or no change. Alternatively, in subregions of the hippocampus, either potentiation, depression or no change could dominate. In this second case, depending on the subregion sampled, each electrode will report only one type of those changes. In accordance with this second scenario, a previous study (Whitlock et al., 2006) that sampled fear-learning related changes at different locations over the dorsal CA1 found that some locations showed potentiation, some showed depression while the rest showed no change. Consequently, averaging the changes across the different locations cancels out synaptic strength changes and results in little to no average effect of learning (Whitlock et al., 2006). Therefore, to identify learning effects, we examined potentiation and depression separately by selecting respectively, the most and least potentiated electrodes within each mouse.

**Figure 2.**
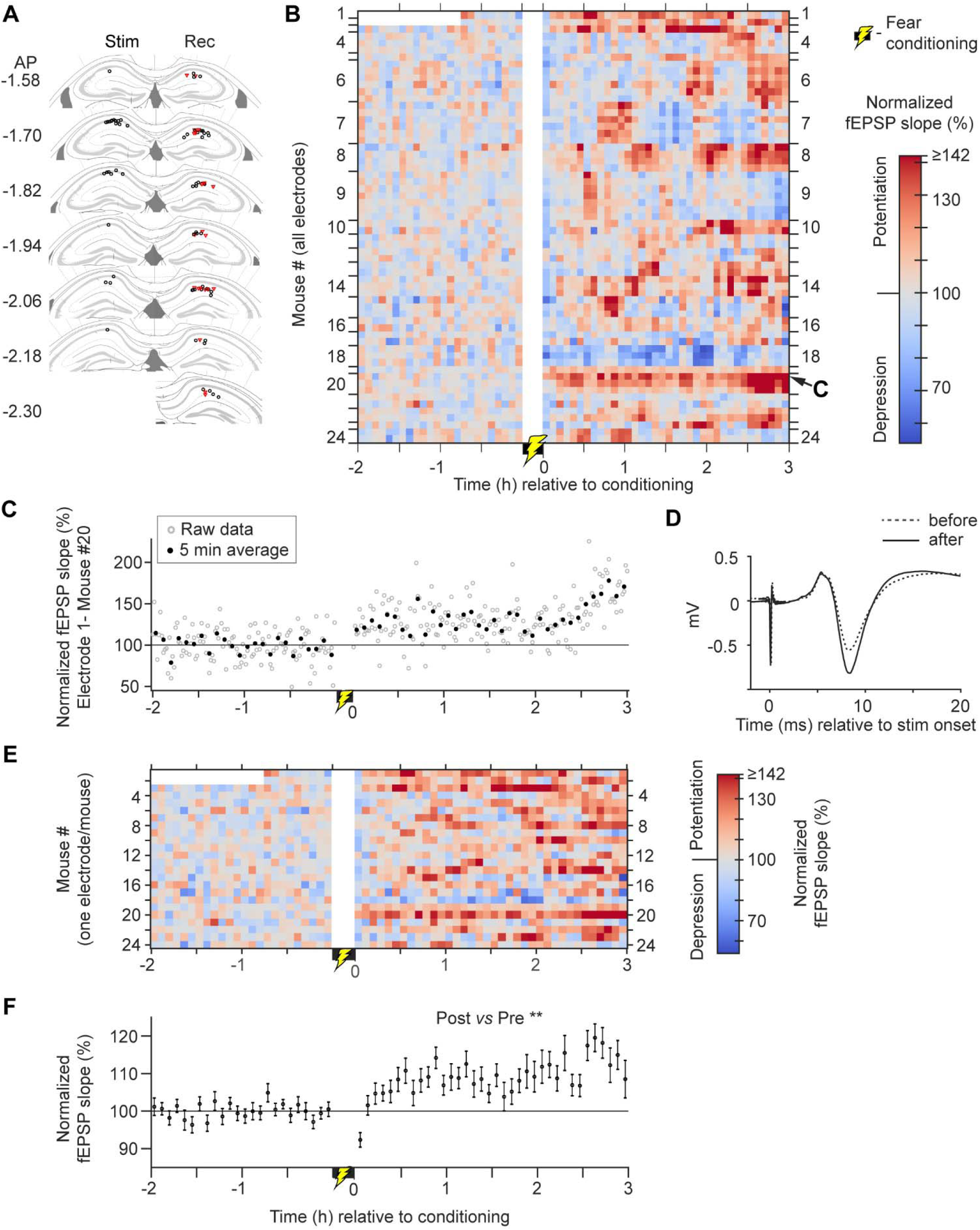
In vivo synaptic strength changes associated with cFC. ***A,*** The locations of the tips of the stimulating (stim) and recording (rec) electrodes were mapped onto a standard mouse atlas (Franklin and Paxinos, 2008). Red triangles indicate the single most potentiated electrode out of each mouse, and those electrodes were used to indicate the largest measurable potentiation and were used in panels ***E*** & ***F***. ***B***, A heat map of the slopes of the fEPSP responses normalized to the baseline. Each row shows data from a single electrode. Rows between the adjacent ticks on the vertical axis indicate the set of electrodes recorded from the same mouse. The foot-shock conditioning period is marked by the yellow lightning bolt symbol near time zero. Mice #1 & 2 had only 45 min of baseline data. Mice # reflects the ascending chronological order of the recording dates. ***C***, The normalized fEPSP slopes from the most potentiated electrode of the set used to record from mouse #20 in ***B***. The gray data points are the raw data, and the black points represent the 5-minute average of the raw data. ***D,*** Stimulation response data from ***C*** averaged across all stimulation repetitions before (dotted trace) and after (continuous trace) fear conditioning. ***E,*** A heat map of the normalized fEPSP slopes from the most potentiated electrode set (red triangles in ***A***). ***F,*** The across-mouse average of the 5-min binned normalized slopes shown in ***E***. Error bars show SEM, and ** p < 0.01, is the significant increase in fEPSP slopes in the post-conditioning period relative to baseline.

To quantify fear conditioning-related synaptic strength changes, we first normalized all fEPSP slopes of a given electrode to the mean fEPSP slope of the baseline period of the same electrode. Figure 2B shows the normalized fEPSP slopes of all the electrodes. There were multiple recording electrodes (2-7) in most mice. For potentiation, we chose one electrode from each mouse as follows. In the subset of mice (18/24) that had multiple electrodes, for each mouse, we chose a single electrode that had the largest potentiation. To this set of selected electrodes, we added the electrodes from the rest of the mice (6/24) that had a single electrode per mouse. Note that although depression or no change may be found in some of the electrodes of the single-electrode mice, to get a conservative estimate of potentiation, we decided to combine all the single electrode cases with the data from the most-potentiated electrodes from mice that had multiple electrodes. This process will lead only to an underestimation of maximum measurable synaptic potentiation.

Following fear conditioning, visual inspection indicates that many electrodes showed a small potentiation, some showed no change, and depression was rare (Figure 2B). Data from the most-potentiated electrode of the implanted set of an example mouse are shown in Figure 2C where a sustained increase in synaptic strength can be seen, with the averaged fEPSP responses shown in Figure 2D. Data from all of the most-potentiated electrodes are shown in Figure 2E with warmer red colors indicating potentiation. On average, the post-conditioning period showed substantial potentiation compared to baseline (Figure 2F). To assess the statistical significance of the results, we used a linear mixed model (LMM) approach, treating the sequence of fEPSP slopes as an interrupted time series (Wagner et al., 2002), where the changes in post-conditioning fEPSP slopes result from fear conditioning “interrupting” the sequence of baseline fEPSP slopes (see Methods). The analysis showed that the baseline was stable for 2h, i.e., there was no significant change of fEPSP slopes over time: F(1, 1544.1) = 0.21, p = 0.651, LMM. Following cFC learning, there was a significant increase in the fEPSP slopes: F(1, 1618.8) = 11.98, p = 0.001, LMM. In the post-conditioning period, the synaptic strength showed a tendency to grow larger with time (F(1, 1477.7) = 3.6, p = 0.059, LMM), suggesting that additional factors may come into play as time progresses.

Next, instead of using a single electrode from each mouse, we averaged all the electrodes within each mouse and performed the same above analysis. The results showed a significant potentiation associated with learning (Supplemental Figure S1A-B), suggesting that most of the sampled subregions of the hippocampus exhibited potentiation: i.e., a significant increase in the fEPSP slopes following learning (F(1, 1417.6) = 5.8, p = 0.016); stable baseline (F(1, 1303.6) = 0.435, p = 0.51). In the post-conditioning period, there was no significant growth of the synaptic strength over time: F(1, 1219.8) = 2.63, p = 0.105, LMM. These data suggest that after fear conditioning, a subset of commissural synapses undergo long-term potentiation in the dorsal CA1 region.

Next, we examined if changes in the behavioral state and animal motion could have contributed to the increased synaptic strength in the post-conditioning period. Associative and commissural synaptic inputs from CA3 have a stronger excitatory effect on CA1 fEPSP during NREM sleep, compared to the awake state or REM sleep (Segal, 1978; Winson & Abzug, 1977, 1978a, 1978b). Although all our experiments were done during the awake phase of the light/dark cycle, we still considered the possibility that after fear conditioning, the mice may sleep, which could produce increased fEPSPs that would inflate the appearance of synaptic potentiation. In fact, such NREM sleep-mediated enhancement of synaptic strength could potentially explain the noticeable increase of fEPSP slopes during the final 30-minute period of post-conditioning recording (Figure 3A, downward arrow) from the same example mouse shown in Figure 2C. Others have shown that NREM sleep can be detected in the field recordings by a drop in the theta frequency band (i.e., 5-11 Hz) power and an increase in the delta frequency band (1-4Hz) power (Grosmark et al., 2012; Mizuseki et al., 2011; Rothschild et al., 2017; Sirota et al., 2008). That change is indeed the case during the 30-minute time period (Figure 3B, labelled NREM) in the example mouse. Following previously published guidelines (Grosmark et al., 2012), we identified sleep states using the theta/delta ratio and the motion index (see Methods, Supplemental Figure S2). Time segments with a sustained decrease in the theta/delta ratio (Figure 3C) and corresponding immobility (Figure 3D) were labeled as NREM states. REM states were identified by an increase in the theta/delta ratio that corresponded with animal immobility (Supplemental Figure S3). During awake periods, the motion level was variable, arising from multiple awake states (Hulse et al., 2017), and we grouped them into one “awake” behavioral state. We then averaged the waveform of all the stimulated fEPSPs within awake and NREM states. In the awake state, the average stimulation response after cFC was higher than that of baseline (Figure 3E). Interestingly, during post-conditioning NREM periods, this effect was further enhanced, possibly reflecting participation of NREM in memory-consolidation mechanisms. To explore this further, we examined the NREM effect across the mouse population.

**Figure 3.**
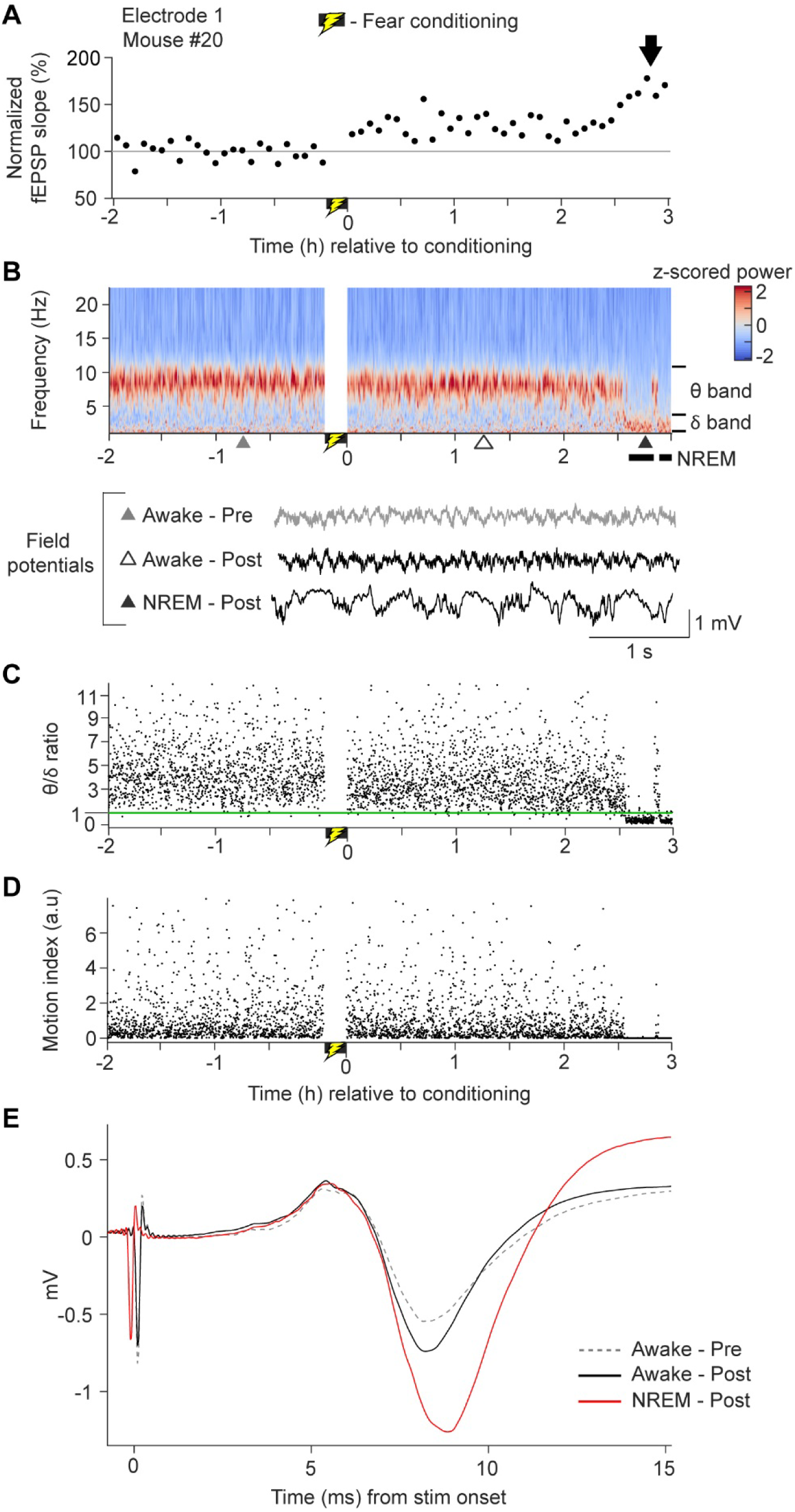
An example of NREM sleep associated with fear conditioning, and NREM effects on synaptic strength. The foot-shock conditioning period is marked by the yellow lightning bolt symbol near time zero. ***A,*** Normalized 5-min binned fEPSP slopes from a single electrode from mouse #20 as in Figure 2B-C. The downward black arrow indicates an abrupt increase in synaptic potentiation in the last 30 min. ***B,*** Top, Spectrogram (z-scored across frequencies) of continuous broadband data from the electrode in ***A***. Black bars below the x-axis indicate the NREM periods. Theta frequency (θ, 5-11 Hz) and delta frequency (δ, 1-4 Hz) band boundaries are shown on the right. Bottom, Ra w field potential traces of 4 s duration taken at time points marked by triangles at the bottom of the spectrogram. Scale bars at the bottom right apply to all three traces. ***C***, Ratio of power in the theta band over the delta band (θ / δ computed in 5-s contiguous time windows. ***D***, Motion index computed in 5-s contiguous time windows in arbitrary units (a.u). ***E*,** Average fEPSP responses during baseline (dotted trace), post-conditioning awake period (black trace), and post-conditioning NREM sleep period (red trace).

Sleep analysis was restricted to the 5 h recording period (2h pre- and 3h post-conditioning). NREM sleep occupied most of the total sleep period (NREM: 95%, REM: 5%, Supplemental Figure S3). The median length of the NREM state was 145 s (IQR: 75 – 263 s, n = 195 NREM segments from 20 mice) and that of REM state was 74 s (IQR: 57 – 95 s, n = 25 REM segments from 12 mice). Figure 4A shows the NREM and REM states along with the normalized fEPSP slopes. Visual inspection of Figure 4A shows that only a few animals slept briefly during the baseline period (median 0%, min 0%, max 16%, Figure 4B). However, a majority of animals slept intermittently after fear conditioning (median 13%, min 0%, max 36%, Figure 4B), resulting in a significantly higher proportion of time spent sleeping in the post-conditioning period (Figure 4B): p = 1.9 x 10^-6^, n = 24, sign test. As seen in the example mouse data (Figure 3), the averaged fEPSP slope during the NREM epochs was higher than that of the awake state for each mouse (Figure 4C): n = 19, where mice with less than a total of 5 stimulus pulses within each behavioral state were excluded. On average, the NREM state slopes were 1.29 ± 0.014 times higher than those of awake state (Figure 4D): p = 8.3 x 10^-15^, n = 19, ratio t-test. Of note, the NREM-mediated enhancement of fEPSP slopes was also evident during the baseline in three mice (Figure 4C-E, filled magenta circles). This suggests that NREM state can boost the fEPSP slopes irrespective of cFC learning, which is in accordance with previous studies (Winson & Abzug, 1977, 1978a, 1978b) that also found similar enhanced synaptic responses during NREM state in non-learning conditions.

**Figure 4.**
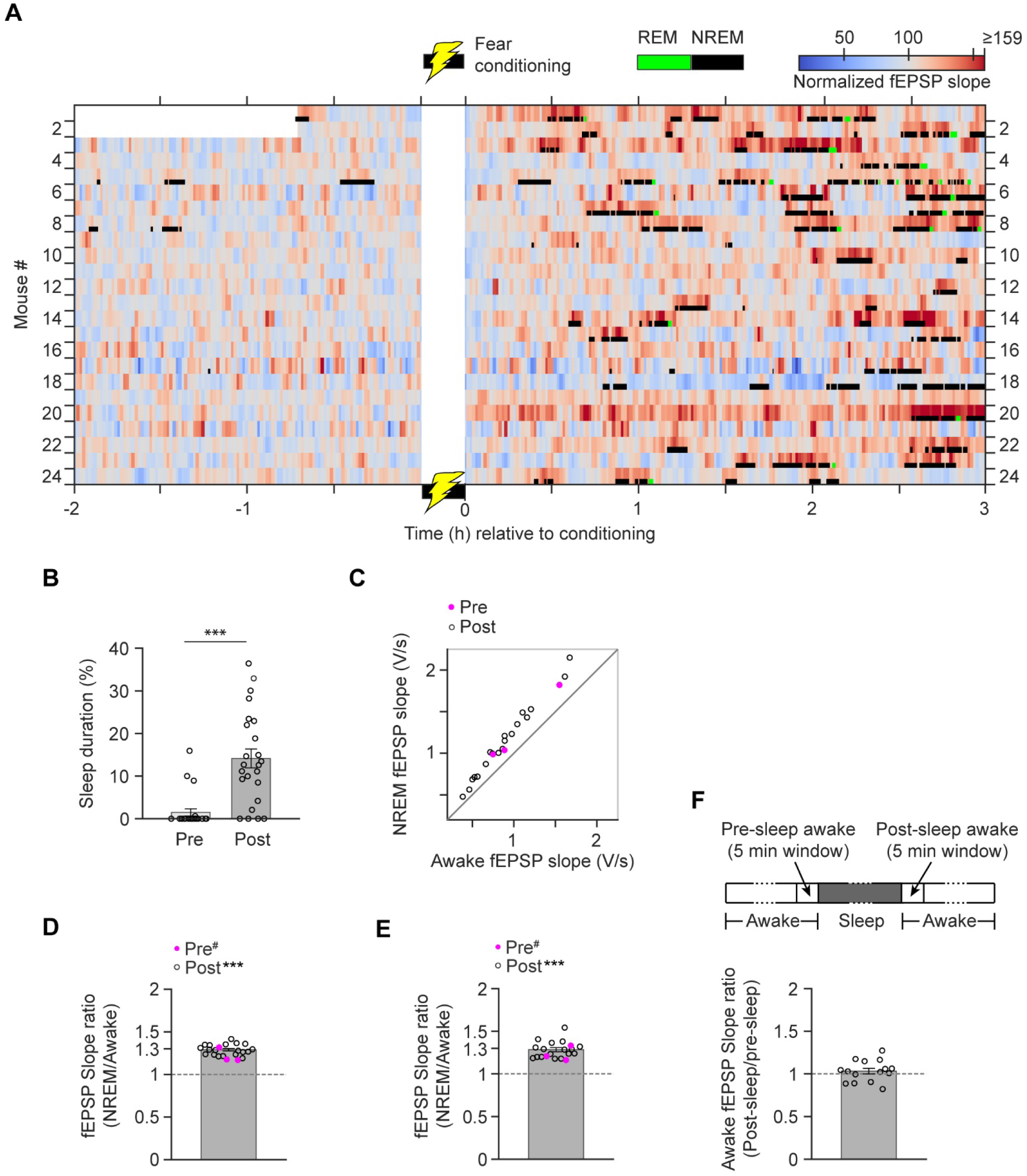
cFC associated changes in sleep and the effect of NREM sleep on synaptic strength. The foot-shock conditioning period is marked by the yellow lightning bolt symbol near time zero. ***A,*** Baseline-normalized fEPSP slopes (non-binned slopes, smoothed with a Gaussian kernel of 45s standard deviation) overlaid with identified REM (green) and NREM (black) periods. Each row corresponds to data from a maximally potentiated single electrode from each mouse (the same electrodes as in Figure 2E). ***B,*** Percentage of time spent sleeping (REM+NREM) during the baseline (Pre) and post-conditioning (Post) periods. ***C,*** Scatter plot of NREM versus awake non-normalized, average fEPSP slopes (Volts/s, with the sign reversed). Each data point represents a single electrode in panel ***A***. Open circles represent the post-conditioning period (n = 19). Filled magenta circles represent the baseline period (n = 3). ***D,*** Data from panel ***C*** are plotted as the ratio of the NREM period fEPSP slopes over the awake period slopes. Mean and error bars are computed with post-conditioning data only; *** p <0.001, for the post-conditioning period, slope ratio is significantly different from 1; #, indicates no statistics were done on the baseline sleep due to the small sample size. ***E,*** As in ***D*** except that the NREM-to-awake ratio was computed by considering only the awake periods that immediately follow each NREM segment. ***F,*** Top: Schematics (not drawn to scale) showing the locations of awake state (white) 5-min time windows (indicated by arrows) in relation to a sleep episode (gray). Dotted lines denote variable length of epochs. Bottom: Ratio of average fEPSP slope of post-sleep awake window (Post-sleep awake) over that of the pre-sleep awake window (Pre-sleep awake); n = 15.

Next, we asked if the NREM-mediated enhancement of synaptic strength outlasts the sleep episode that contains the NREM epochs. Sleep episodes are naturally interrupted by brief awakenings (Chemelli et al., 1999; Franken et al., 1999; Huang et al., 2006; Léna et al., 2004; Lima et al., 2017; Perez-Atencio et al., 2018; Watson et al., 2016). Accordingly, we grouped a set of NREM and REM segments interspersed by brief awakenings (< 1.5 min) into a single sleep episode (Supplemental Figure S2-S3). If the NREM-mediated synaptic enhancement continued into the adjacent awake state, then the NREM-to-awake state fEPSP slope ratio computed using adjacent sleep-awake segment pairs, should not be different from 1. This continuation of the NREM enhancement is not what we found. The across-mouse NREM/awake ratio average was 1.28 ± 0.024, which was significantly different from 1 (Figure 4E): n = 17, p = 2.42 x 10^-10^, ratio t-test. Since any extension of the NREM effect into the awake state may dissipate over time, to favor detection of any lasting effect of NREM, we shortened the roughly 26 min awake period of the above analysis to a maximum of 15 min post-sleep. The results remained the same, with NREM/awake ratio = 1.26 ± 0.025, which is again significantly different from 1: n = 17; p = 2.26 x 10^-9^, ratio t-test. This result suggests that NREM-mediated enhancement of synaptic strength does not outlast (intact) the sleep episode on the 15-min time scale.

Although the NREM effect did not outlast the sleep episode, the NREM state, and sleep in general could renormalize synaptic strength (Norimoto et al., 2018; Tononi & Cirelli, 2020; Vyazovskiy et al., 2008). We tested if renormalization occurs (on the time scale we are looking) in commissural synapses by asking if the synaptic strength is lower in the awake state that follows a sleep episode compared with that measured in the awake state preceding the sleep episode as previously demonstrated in the cortex (Vyazovskiy et al., 2008). In the post-conditioning sessions, we selected two narrow 5-min awake-state time windows - one that immediately precedes (pre-sleep) and another that immediately follows (post-sleep) a given sleep episode (Figure 4F, Top). For the electrode set in Figure 4A, we computed the average fEPSP slope within the post- and pre-sleep awake state windows and obtained the post/pre ratio of fEPSP slope. The mean length of the selected sleep episodes was 13.7 ± 1.2 min. The mean length of NREM state within these sleep episodes was 11.3 ± 0.9 min. The level of animal movement during the pre- and post-sleep 5-min awake windows were comparable: average instantaneous animal speed at the time of the synaptic strength measurements with pre-sleep = 4.0 ± 0.4 cm/s, post-sleep = 4.7 ± 0.5 cm/s. There was no significant difference between pre- and post-sleep speeds: p = 0.25, n = 15, paired t-test. We found (Figure 4F) that the mean post-sleep/pre-sleep awake window synaptic strength ratio was 1.03 ± 0.03, which is not significantly different from 1: p = 0.43, n = 15, ratio t-test. This suggests that brief intervening sleep periods do not alter pre-existing awake state synaptic strength.

In addition to the effect of NREM sleep on fEPSP size, the speed of animal motion during the awake state has been shown to be negatively correlated with fEPSP size (Kemere et al., 2013). To characterize the effect of animal motion on synaptic strength, we computed the correlation between the normalized fEPSP slope and the instantaneous speed of the mouse. Since there may be a time delay for the animal’s movement to impact the synaptic strength, we analyzed movement speed at the onset time of the stimulation pulse (i.e., zero time lag) and at various time points before (i.e., negative lags) and after (i.e., positive lags) the stimulation pulse onset. Note that to get a large sample size and to avoid sleep and learning-related changes influencing the fEPSP slope-speed correlation, we only used the 2h baseline data from all mice (n = 36) in our dataset with any rarely present sleep episodes excluded. Since each mouse may naturally move slower or faster, each could have a different speed distribution. Hence, we computed a repeated measures correlation (RMC) (Bakdash & Marusich, 2017), which computes a single correlation coefficient for the whole mouse population, taking into account any mouse-specific distribution of speed and fEPSP slope. We found a significant negative correlation: r = −0.18, p < 10^-10^, Bootstrap test with a 95% Bootstrap confidence interval (−0.21, - 0.14) between normalized fEPSP slope and instantaneous speed measured at the time of the fEPSP measurement (zero lag, Figure 5A). Note that the linear model was fit simultaneously to all 36 mice. In Figure 5A, to represent the entire span of the mouse speeds, we chose mice that had the lowest, median and highest average speed. To examine if the speed of the animal measured before or after the stimulation pulse is correlated with fEPSP slope, we computed the RMC at a range of positive and negative lags (Figure 5B). We found the maximum strength of correlation near zero lag suggesting that the effect of speed on synaptic strength, if causal, is rather immediate.

**Figure 5.**
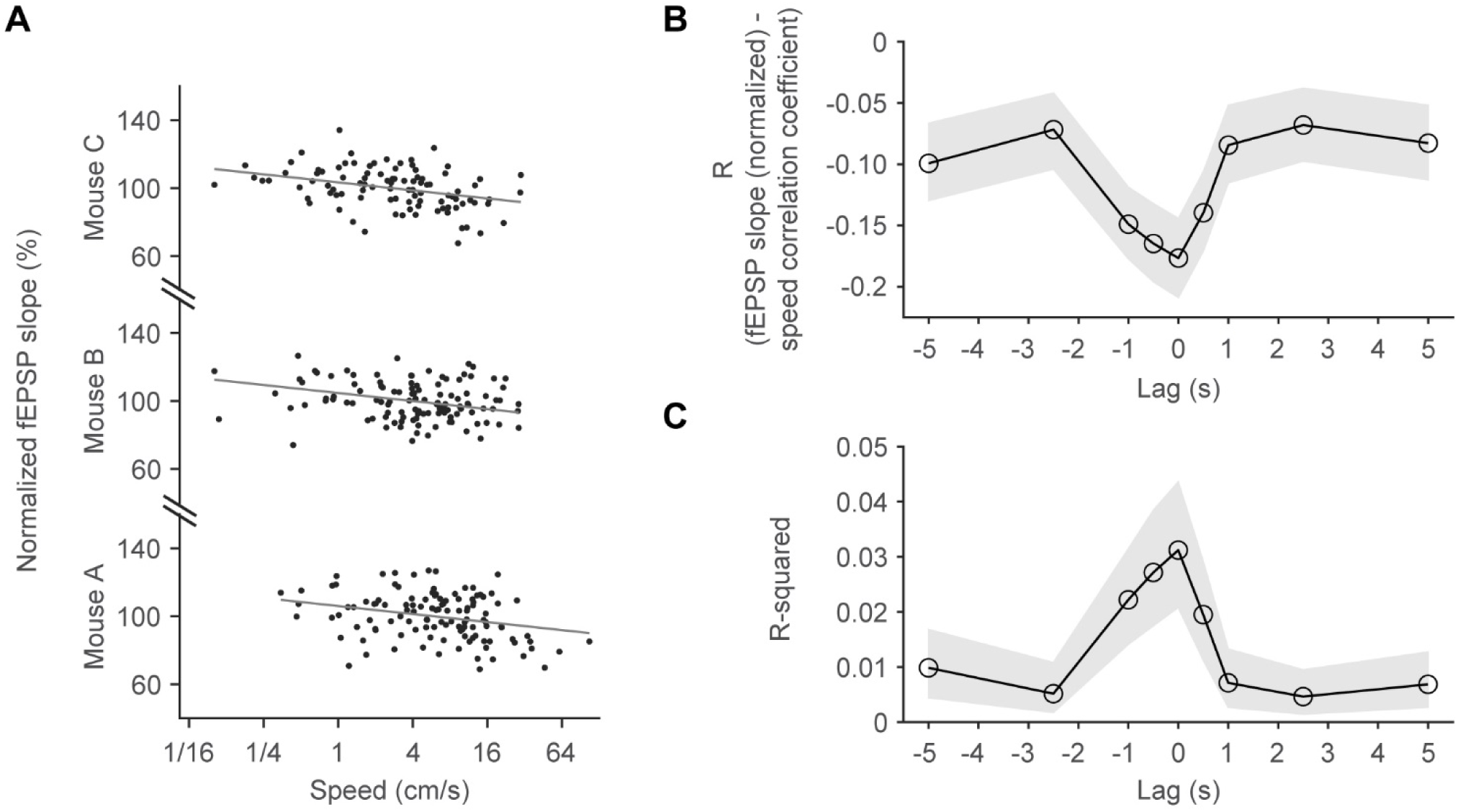
Effect of speed of animals’ movement on synaptic strength. ***A,*** Normalized fEPSP slopes of the baseline period, which is plotted as a function of the instantaneous speed that was measured at the time of stimulus pulses. Data are shown for three mice that represent the lowest (Mouse C), median (Mouse B) and highest (Mouse A) average mouse speed across the population (n = 36; 24 from shocked group, 12 from control group; see Figure 7). The regression lines (gray) represent a global (n = 36) linear model fit that assigns a single gain (slope of the line) and mouse-specific intercepts. ***B,*** Repeated-measures correlation coefficient (R, from the model fit in A) computed using instantaneous speeds measured at different time delays (lags) from the stimulus pulse onset time, which is time zero. ***C,*** R-squared values computed based on data in ***B***. In ***B*** and ***C,*** the gray regions represent 95% bootstrap-based confidence intervals.

Although speed is significantly correlated with fEPSP slope, this correlation explained only a small amount (∼3%) of the variance in fEPSP slope: r^2^ = 0.031 (Figure 5C). However, after cFC, animals may move systematically slower relative to their speed during the baseline. Because speed is negatively correlated with fEPSP size, such a reduction in speed will inflate the learning-related effects seen in the post-conditioning period. Hence, to control for the speed effect, we included the instantaneous speed (measured at zero lag) as a covariate in all our linear mixed model analyses (see Methods and Supplemental materials). Also, evidence implicates sleep as an integral component of memory consolidation (Abel et al., 2013; Klinzing et al., 2019; Walker & Stickgold, 2006). Hence, the sleep periods following cFC may serve important functions related to memory consolidation. However, for a conservative estimate of the synaptic potentiation that might contribute to learning, we set out to remove the slope data corresponding to sleep episodes that would exaggerate the synaptic potentiation. Note that we did not have enough stimulation pulses within REM sleep to examine its separate effect on fEPSP slopes. However previous work has shown that NREM and REM have different effects on fEPSP slopes (Winson & Abzug, 1978b). Since our goal was to have the awake state for the pre-versus post-conditioning comparisons, we removed sleep episodes (which included both REM and NREM) leaving us with only the awake state.

Starting with the original complete dataset (Figure 2B), we removed the fEPSP slopes corresponding to the sleep episodes (Supplemental Figure S2-S3) from baseline and post-conditioning periods and then normalized all slopes to the baseline. As described for Figure 2, a single electrode was then chosen from each mouse to form the most-potentiated electrode set. The average of the fEPSP slopes after removing the sleep segments still shows potentiation (Figure 6A). Data from the same electrodes are shown in Figures 2E and 6A to aid in comparison of the synaptic potentiation with sleep (Figure 2E) and without sleep (Figure 6A). Interestingly, even after removing the NREM-mediated enhancement of synaptic transmission and controlling for the speed effect, the fEPSP slopes were still significantly increased in the post-conditioning period: F(1, 1623.6) = 11.8, p = 0.001, LMM. Within the baseline and post-conditioning period, fEPSP slopes did not change over time: F(1, 1556.3) = 0.005, p = 0.95 for the baseline; F(1, 1524.1) = 0.002, p = 0.97, for the post-conditioning period, LMM. The result indicates that the baseline remained stable and that the synaptic potentiation effect persisted throughout the post-conditioning period. The effect of speed was significant (F(1,5880) = 242.2, p < 0.001), suggesting that it contributes to the measured synaptic strength size. In a previous study (Zhou et al., 2017), they used a different fear conditioning paradigm in which the day before conditioning the animals were pre-exposed to the conditioning and a novel spatial context. In the 1h post-conditioning period that was examined, that study did not find any learning effect. Restricting our post-conditioning data to the same 1h period still yielded a significant learning effect, F(1,1246.2) = 9.5, p = 0.002; significant effect of speed, F(1,3919.2) = 133.6, p < 0.001; non-significant effect of time in baseline, F(1,1156.8) = 0.004, p = 0.949 and post-conditioning period, F(1, 1187.3) = 0.1, p = 0.75. The difference between the Zhou et al. (Zhou et al., 2017) study and ours suggests that the commissural pathway plasticity depends on task requirements.

**Figure 6.**
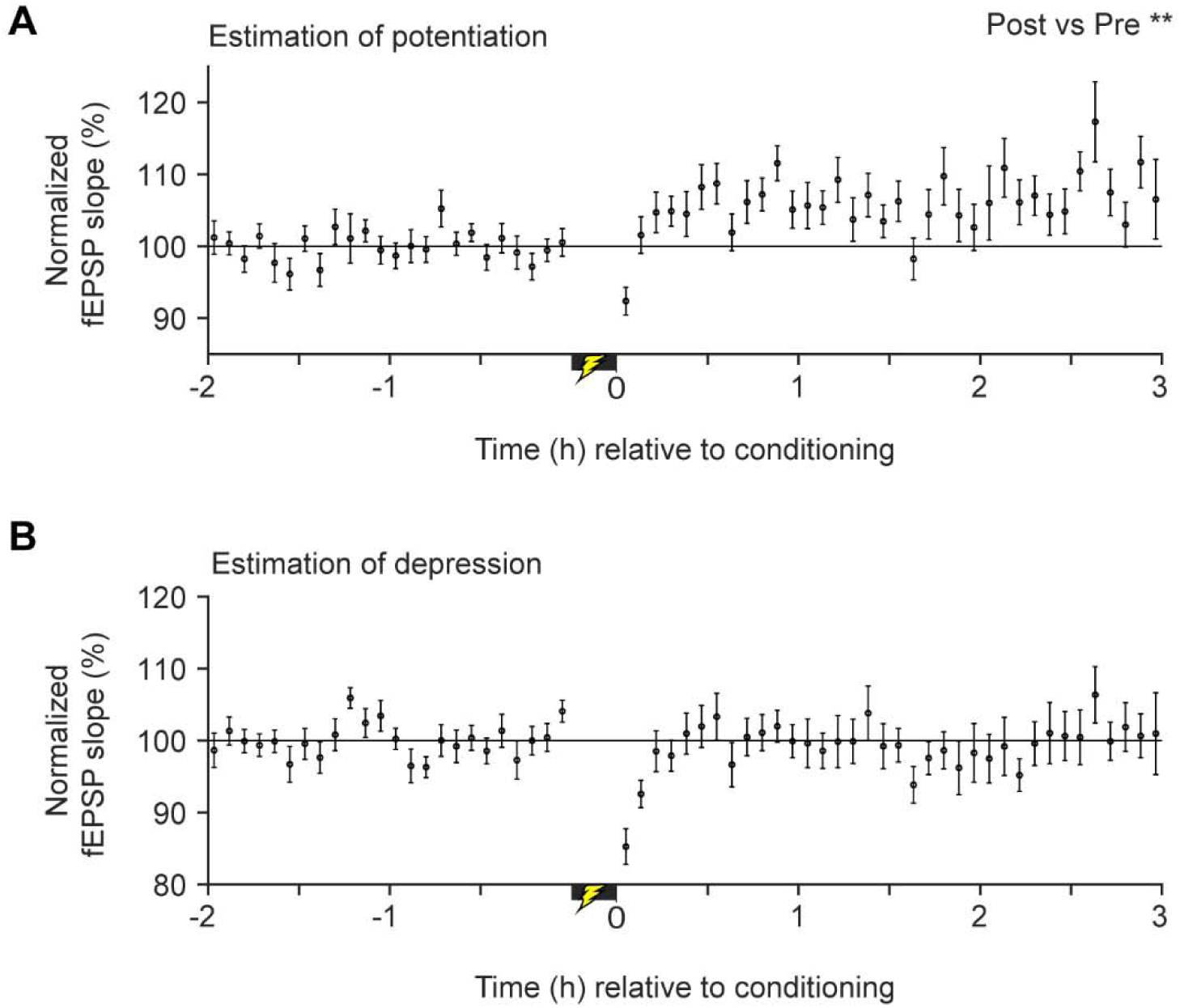
In vivo synaptic strength modifications associated with cFC after taking sleep into account. ***A,*** Average normalized fEPSP slopes from the most potentiated electrodes (n = 24, 1 electrode/mouse) after removing the fEPSP slope data corresponding to sleep episodes (Supplemental Figure S3). ** p < 0.01, significant increase in fEPSP slopes in the post-conditioning period relative to baseline. ***B***, as in ***A*** except that least potentiated electrodes (n = 18, 1 electrode/mouse) were used. Error bars: SEM.

Next, instead of using a single electrode from each mouse (as in Figure 6A), we averaged all the electrodes for each mouse and examined the synaptic potentiation of the cFC learning. The results remained the same (Supplemental Figure S1C). There was still a significant effect of learning, F(1,1443.1) = 4.25, p = 0.039, and a significant effect of speed, F(1,5618.2) = 402, p < 0.001; but there was not a significant effect of time in the baseline, F(1,1352.3) = 0.067, p = 0.8 or in the post-conditioning period, F(1, 1309.8) = 0.001, p = 0.97. These results suggest that the fear conditioning resulted in long-term potentiation of the commissural pathway in a subset of locations in the dorsal CA1 region and that this effect cannot be explained by the sleep state or speed-related covariations during the awake state. Although fear conditioning results in long-term potentiation at the population level, the magnitude of potentiation did not correlate with the behavioral measure of memory (Supplemental Figure S4): F(1,22) = 1.21, p = 0.28, linear regression. The magnitude of potentiation explained only a small amount (∼5%) of the variance in the freezing level: r^2^ =0.052. This result suggests that the relationship between commissural pathway potentiation magnitude and freezing level may be more complex than a simple linear one or that our electrode placements sampled different proportions of the overall synaptic potentiation associated with the cFC-induced learning.

Long-term depression has also been proposed to participate in learning and memory (Bear, 1996; Kemp & Manahan-Vaughan, 2004; Malenka & Bear, 2004). To test whether cFC resulted in any significant decrease in the average fEPSP slopes, we first removed slope data corresponding to the sleep episodes and normalized the slopes to the baseline as we did in Figure 6A. Then, from each mouse that had multiple electrodes (n = 18 mice), the least-potentiated electrode was selected. Unlike our analysis of potentiation, mice with single electrodes (n = 6) were excluded as any potentiation in some of them would mask any possible depression effects. To control for speed effects, we also included the instantaneous speed as a covariate in our linear mixed model. Following cFC, relative to baseline, there was no significant change in fEPSP slopes (Figure 6B): F(1, 1239.1) = 1.29, p = 0.256, LMM. Within the baseline and post-conditioning period, the fEPSP slopes did not change significantly over time: baseline, F(1, 1187) = 0.006, p = 0.94; post-conditioning period, F(1, 1160) = 0.023, p = 0.88, LMM. These results indicate that the baseline remained stable and that the synaptic strength did not undergo depression, but rather, synaptic strength remained unchanged throughout the post-conditioning period. These results show that fear conditioning under our protocol did not result in measurable long-term depression of the commissural synapses onto the dorsal CA1 region.

Since we did not habituate the animals to the fear conditioning chamber and the mice had not been used in any previous behavioral tasks, our fear conditioning task inherently included exploration of a novel place (i.e., the conditioning chamber). Exploration of a novel place may have itself contributed to the synaptic potentiation we observed. To address this issue, we tested synaptic strength before and after a sham-fear conditioning procedure where the animals explored the conditioning chamber, but no foot-shocks were given. Figure 7A shows the location of the electrode tips. The quality of the fEPSP measurement in the shocked and control conditions were comparable. The depths of the recording electrodes were not significantly different from those of the shocked group: using all the electrode sets, n = 23, 62 (control, shocked), p = 0.38, t-test; using the maximally potentiated electrode set, n = 12, 24 (control, shocked), p = 0.337, t-test. There was no significant difference in the stimulation current intensity used for the two conditions: 29.0 ± 3.2 µA for the control, 33.2 ± 1.5 µA for the shocked, n = 12, 24 (control, shocked), p = 0.187, t-test. There was no significant difference in the size of the fEPSP response as indicated by the raw slope value of the fEPSP response between the two conditions using all the electrode-sets: −0.73 ± 0.07 V/s (control), −0.79 ± 0.04 V/s (shocked), n = 23, 62 (control, shocked), p = 0.45 by t-test; using the maximally potentiated electrode set, −0.75 ± 0.11 V/s (control), −0.82 ± 0.06 V/s (shocked), n = 12, 24 (control, shocked), p = 0.57 by t-test.

**Figure 7.**
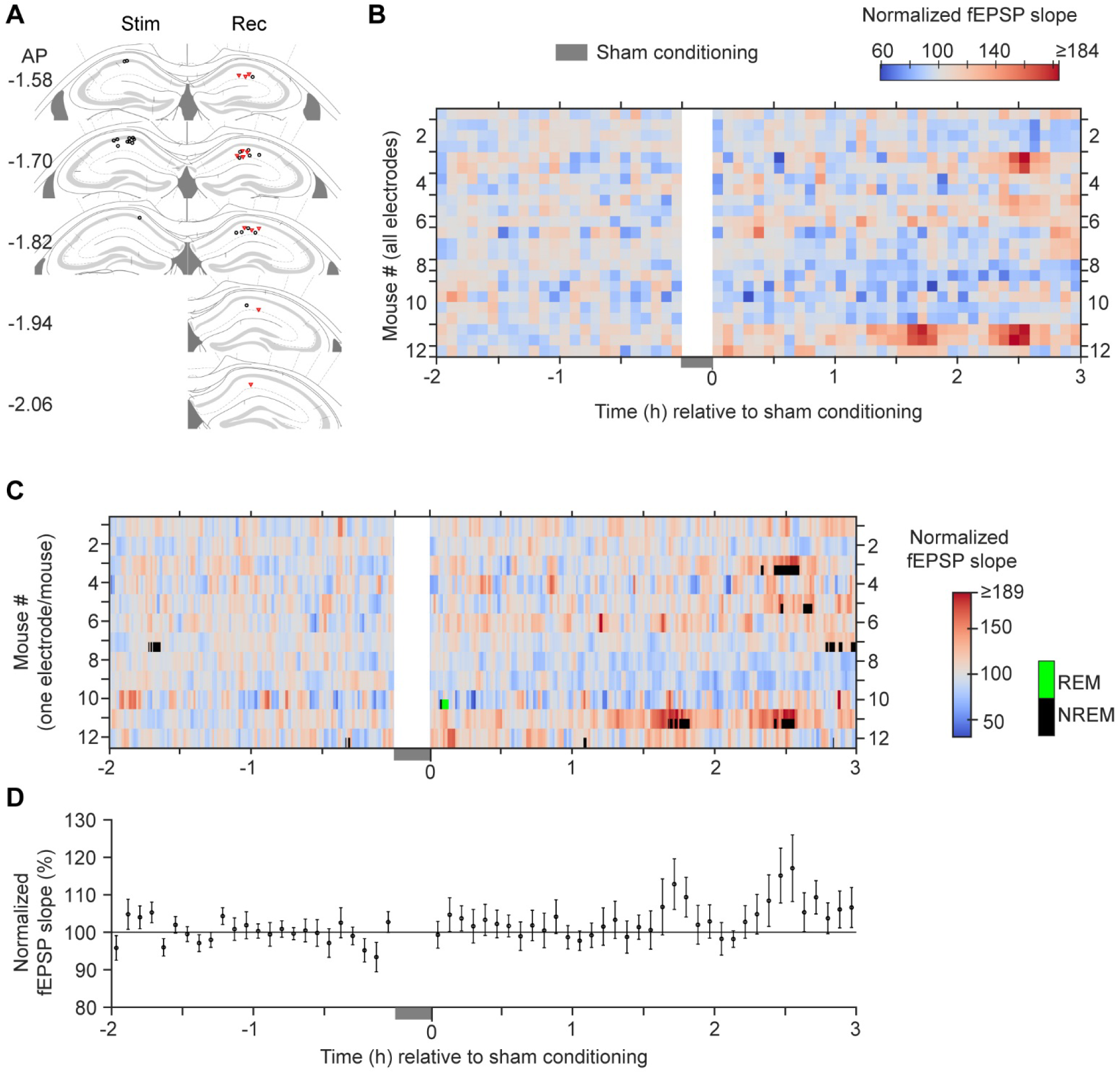
In vivo synaptic strength modifications associated with sham-conditioning. ***A,*** Locations of the tips of the stimulating (stim) and recording (rec) electrodes mapped onto a standard mouse atlas (Franklin and Paxinos, 2008). Red triangles indicate the single most potentiated electrode out of each mouse. Those electrodes were used to indicate the largest measurable potentiation and were used in panel ***C-D. B***, Heat map of the baseline-normalized fEPSP slopes. Each row shows 5-min binned data from a single electrode. Rows between the adjacent ticks on the vertical axis indicate the set of electrodes recorded from the same mouse. The sham-conditioning period is marked by a gray box near time zero on the x-axis. ***C***, Baseline-normalized fEPSP slopes (non-binned slopes, smoothed with a Gaussian kernel of 45s standard deviation) overlaid with identified REM (green) and NREM (black) periods. Each row corresponds to data from a maximally potentiated single electrode from each mouse (n = 12, 1 electrode/mouse). ***D,*** Average normalized fEPSP slopes from the most potentiated electrodes (shown in ***C***) after removing the fEPSP slope data corresponding to sleep segments (Supplemental Figure S5). Error bars are SEM.

The normalized fEPSP slopes from all mice (Figure 7B) shows that there was very little change in synaptic strength following the sham conditioning. The sleep patterns (Figure 7C, Supplemental Figure S5) indicated that most animals stayed awake throughout the duration of the recording (baseline: median, 0%, min, 0%, max, 4%; post-conditioning: median, 0%, min, 0%, max, 8%) with no significant change in the sleep after the sham-fear conditioning procedure compared to baseline (p = 0.219, n = 12, signtest). This sleep pattern is in contrast to that of the shocked group, which slept more during post-conditioning relative to baseline: p = 1.5 x 10^-4^, n = 24, 12 (shocked, control), one-tailed ranksum test on post-pre difference in sleep %. However, similar to the shocked group, when the sham-trained animals went into NREM sleep, the fEPSP responses were higher relative to those of awake state (Figure 7C). For the 4 mice with at least 5 min of NREM sleep, the fEPSP slope NREM/awake ratio was 1.52, 1.26, 1.31 and 1.5. Hence, following the same procedure used for the fear-conditioned animals (Figure 6A), we chose the most potentiated electrodes after removing the fEPSP slope data that corresponded to the sleep episodes. To control for speed effects, we also included the instantaneous speed as a covariate in our linear mixed model. The across-mouse average (Figure 7D) shows that there was no significant change in synaptic strength in the post-conditioning session relative to baseline: F(1, 891) = 1.28, p = 0.259, LMM. The effect of speed was significant: F(1, 3291.3) = 143.1, p < 0.001. Within the baseline and post-conditioning period, fEPSP slopes did not change over time: baseline, F(1, 863.7) = 1.54, p = 0.216; post-conditioning periods, F(1, 838.3) = 3.03, p = 0.082, LMM. These results indicate that the baseline remained stable and that the synaptic strength remained unchanged throughout the post-conditioning period. These results suggest that exploration of a novel place – the fear conditioning chamber – was not sufficient to result in significant measurable changes in synaptic strength. Furthermore, after accounting for the speed effect, relative to the sham-conditioning control, fear-conditioned animals showed a significant potentiation in the post-conditioning session with a significant pre-post x shock condition interaction: F(1, 2361.2) = 4.41, p = 0.036, LMM. Taken together these results clearly demonstrate that fear conditioning potentiates the commissural pathway of the dorsal CA1 and that this potentiation cannot be explained by the sleep state, the animal’s movement speed, or exploration of a novel environment.

## Discussion

Contextual fear-conditioning has been used for decades as a model for associative fear learning. Although numerous studies have observed cFC-related changes at the molecular, cellular, and synaptic levels in the hippocampus (Buffington et al., 2014; Johansen et al., 2011; Nicoll, 2017; Sacchetti et al., 2001; Weng et al., 2018), real-time in vivo synaptic plasticity has not been examined. Our data demonstrate that the dorsal CA1 commissural pathway of the hippocampus undergoes long-term potentiation in vivo in association with our fear conditioning task. Our data also show that sleep is more common after cFC and that the size of the fEPSP responses are bigger during NREM sleep than during the awake state. We also found that the speed of animal movement covaries with the fEPSP size. These observations emphasize the importance of taking an animal’s motion and behavioral state into account when measuring learning-related synaptic changes.

There are numerous commissural fibers connecting the subregions of the two hippocampi in rodents (Laurberg, 1979; Swanson et al., 1978), but the function of this pathway is not well characterized. A study in rats suggests that the dorsal hippocampal CA1 to CA1 commissural pathway is involved in a rapid form (< 24h) of fear generalization (Zhou et al., 2017) that is distinct from previously studied fear generalization that develops over many days (Cullen et al., 2015; dos Santos Corrêa et al., 2019; Pedraza et al., 2016; Wiltgen & Silva, 2007; Wiltgen et al., 2010). However, the rapid generalization that was observed (Zhou et al., 2017) depended on pre-exposure to two different contexts (conditioning and non-conditioning) the day before the conditioning. In this paradigm, contrary to our results, the authors show no plasticity in the commissural pathway in the first roughly 1.5h following the conditioning. However, they did show potentiation 24h post-conditioning when fear generalization is fully developed, suggesting that the potentiation they observed is more related to fear generalization. Given that our experimental paradigm is different from that used by Zhou et al. (2017), the plasticity we observed immediately after conditioning is unlikely to be related to the rapid fear-generalization they describe. Under our experimental conditions, the results suggest that the commissural pathway may be involved in a specific component of the learning task. A potential next step would be to test if the potentiation we observed is related to learning of the conditioning context or related to linking the context with the aversive events occurring in that context. Because exposure to the conditioning chamber (without shocks) did not result in measurable synaptic potentiation, learning the context or place is not sufficient to produce the commissural potentiation that we measured, suggesting the pathway may contribute to linking the aversive event to the context.

For quantifying potentiation and depression, when multiple electrodes were present in our mice, we selected the most- and least-potentiated electrodes respectively. For our experimental settings, this approach was necessary because as reported previously (Whitlock et al., 2006), some locations in the dorsal hippocampus undergo potentiation whereas others undergo depression or no change. Hence averaging across electrodes may mask learning-related plastic changes. Also, our electrode-selection approach did not lead to detecting random noise as potentiation or depression. This is because, in the absence of a learning effect, i.e., in the control groups, potentiation was not detectable either in our study or in the two other previous studies (Whitlock et al., 2006; Broussard et al., 2016). This shows that in the absence of learning, simply picking the largest potentiated electrode does not in itself result in a detectable potentiation. Conversely, even in the shocked group, picking the least potentiated electrodes did not result in significant depression. These data show that any selection bias is below detectable level. More importantly, any sample from one or several electrodes will only randomly measure the synaptic plasticity in that local area, which is not likely to be the area that underwent the most intense potentiation. Had we placed our electrodes in the CA locations that underwent the maximal learning-induced potentiation, the electrodes would have detected a potentiation larger than our reported values, suggesting that it is unlikely that we over-estimated the potentiation. Instead, our results and those from previous related studies (Whitlock et al., 2006; Broussard et al., 2016) suggest that the dorsal CA1 does not undergo identical, uniform potentiation across its whole topological range.

The magnitude of the LTP in the dorsal CA1 we observed (i.e., a roughly 10% change) is smaller than electrical stimulus-induced in vivo LTP, which typically produces a 50-100% change (Martin et al., 2019). The magnitude we observed is comparable to the in vivo plasticity seen in studies of inhibitory avoidance (Broussard et al., 2016; Whitlock et al., 2006) and eyeblink conditioning (Gruart et al., 2006). This smaller in vivo LTP likely results because plastic changes may happen only at a subset of synapses. Given that our fEPSP measurement sums the effect of a large number of synapses most of which are likely unmodified, the contribution of the smaller number of modified synapses in any given recording location gets diluted within the overall average, giving rise to smaller LTP measurements. In addition, the magnitude of the LTP did not correlate with the behavioral measure of memory we employed (freezing). This result is likely due to our electrodes not sampling enough of the locations where plastic changes occurred. If all the electrodes had been placed optimally with respect to where synaptic plasticity occurs (“hits”), the correlation would have been stronger. However, in some mice, large plastic changes may have occurred at one location in correlation with high freezing levels, but our electrodes may have been at a different location that did not undergo plastic changes (“misses”). Such “misses” when present in sufficient numbers can mask the correlation because in these cases, higher freezing levels get associated with lower or no potentiation. Hence, when “hits” and “misses” are pooled, as is likely the case in our dataset, the correlation strength becomes too weak to be significant. Alternatively, the changes in other pathways within the hippocampus (e.g. associative pathways) are more correlative than changes in the commissural pathway we examined. Finally, the relationship between the physiological correlate of memory (CA1 commissural plasticity) and the behavioral correlate of memory (freezing) may be more complex than a simple linear one.

We found a significant negative correlation between movement speed and fEPSP slope in mice as has been found in rats (Kemere et al., 2013). Also, as in the rat study, the strength of this correlation was maximal when the fEPSP slope was correlated with the speed measured at the time of the slope measurement (time lag = 0). However, the variance in the fEPSP slopes explained by speed was relatively small in our dataset (i.e., 3%) when compared to that of the rat study (18%).This difference could arise from behavioral differences where the rats ran in mazes whereas our mice stayed in their home cage during the recordings. Also, there was a difference in the locations that were stimulated in the two studies: the ventral hippocampal commissure/ contralateral CA3 in the rat study versus the contralateral CA1 in our study.

Although all our experiments were done during the dark phase (i.e., awake phase) of the light/dark cycle, we found that the fear-conditioned, shocked mice spent more time in intermittent sleep compared to the sham-conditioned mice (that were not shocked). Although this specific result has not been reported before, there are many studies that have found changes in light-phase sleep architecture in rodents that were subjected to shock conditioning during the light phase or briefly before the light onset (Ambrosini et al., 1993; Bramham et al., 1994; Datta, 2000; Mavanji et al., 2003; Pawlyk et al., 2008; Larry D. Sanford et al., 2003; L. D. Sanford et al., 2003; Smith et al., 1980; Wellman et al., 2008). Similar to our study, a study (Hellman & Abel, 2007) that performed cued fear conditioning 4h into the light phase found an increase in the duration of NREM sleep. However, unlike our study, this increase occurred 12-16h post-conditioning. This comparison suggests that although fear conditioning increases NREM sleep, the type of conditioning procedure (cued versus contextual) and time of conditioning (light versus dark phase) may determine when the NREM sleep changes occur. The mechanisms for the conditioning-induced increased sleep during the awake-phase (i.e., dark-phase) are unclear. Interestingly, the sleep studies that were conducted during the dark phase (Bonnet et al., 1997; del C. Gonzalez et al., 1995; Koehl et al., 2002; Rampin et al., 1991), consistently found that rats that were subjected to brief (∼1h) immobilization stress during the dark phase, slept more in the dark phase compared to non-stressed rats. Thus, foot-shock stress, though different from immobilization stress, can be speculated to contribute to the increased dark-phase sleep following fear-conditioning. Irrespective of the mechanisms behind this increased sleep, sleep disruption for 5h immediately following fear-conditioning leads to memory impairment (Graves et al., 2003). Conversely, inducing sleep immediately after conditioning facilitates memory consolidation (Donlea et al., 2011). During NREM sleep, sharp-wave ripples (which are high frequency oscillations of approximately 200 Hz in the field potential arising from synchronous rhythmic firing of hippocampal neurons) are abundant, and have been implicated in learning and memory (Buzsáki, 2015; Joo & Frank, 2018; Olafsdottir et al., 2018). Ripple disruption during the NREM periods that follow immediately after spatial learning, impairs learning (Girardeau et al., 2009). Together, these studies suggest that the increased sleep we observed likely serves an important function in memory consolidation.

Although the effect of the animal’s brain state on subthreshold activity, spiking, and network activity in the hippocampus have been extensively studied (Buzsáki et al., 1983; Chrobak & Buzsáki, 1996; Hulse et al., 2017; Jarosiewicz et al., 2002; Kay et al., 2016; O’Keefe, 1976; Wilson & McNaughton, 1994), only a handful of rat studies have examined how synaptic transmission is altered during sleep states relative to awake states (Segal, 1978; Winson & Abzug, 1977, 1978a, 1978b). By stimulating the perforant pathway, Winson and Abzug (1977) examined how behavioral states affect synaptic transmission in the ipsilateral monosynaptic (entorhinal cortex, EC, to dentate gyrus, DG), disynaptic (EC to DG to CA3) and trisynaptic pathway (EC to DG to CA3 to CA1). They found that in all three regions (i.e., DG, CA3, and CA1) the population spiking activity was higher during NREM state than during any other brain state. In addition, the fEPSP size of the trisynaptic response in the CA1 region was also higher during NREM. Similarly, the CA1 fEPSP response to activation of the contralateral CA1, was also higher during NREM sleep (Segal, 1978). Our results in mice show that (similar to the rat studies) the commissural CA1 pathway synaptic transmission is enhanced during NREM sleep. In addition, we found that the NREM/awake ratio of fEPSP slopes tightly cluster around a mean value of 1.3 (Figure 4D-E). Thus, irrespective of the size of the awake-period fEPSP responses, in the NREM state, they get scaled by a roughly constant factor. Hence, the aggregate of mechanisms underlying the NREM effect can be conceptualized as a multiplicative gain modulation.

Although the NREM state enhanced synaptic strength, this enhancement was temporary. Once the sleep episode terminated, the synaptic strength returned to the same level observed in the awake period preceding the sleep episode, suggesting that the sleep episode did not renormalize the synaptic strength. This result is in contrast to a previous observation in the cortex (Vyazovskiy et al., 2008) where following a sleep period, the awake state synaptic strength was lower (renormalized) compared to that measured in the awake state preceding the sleep period. Recent studies (Gulati et al., 2017; Norimoto et al., 2018) show that task-related synapses/circuits may be spared from renormalization. As opposed to the previous study (Vyazovskiy et al., 2008) in which the animals did not perform any learning tasks, the cFC learning in our mice could have led to sparing the commissural synapses from renormalization. Another more likely reason for the discrepancy is that the length of our analyzed sleep episodes (∼13 min) is too short for the renormalization to be detectable. The longer sleep periods of ∼ 4h (as in Vyazovskiy et al., 2008) may be necessary for noticeable renormalization.

Considered as an evolutionary precursor to the corpus callosum, the hippocampal commissures have been retained in all mammals (Suárez et al., 2014), indicating their computational importance. Although the functions of hippocampal commissures are largely unknown, one of the proposed functions of the corpus callosum is to integrate lateralized functions across the hemispheres (Aboitiz & Montiel, 2003; Bloom & Hynd, 2005). Given that functional specialization of the left and right hippocampus has been reported in rodents (Jordan, 2020), it is possible that the hippocampal commissures may integrate the lateralized functions recruited during the fear-conditioning task. The plasticity we observed may thus indicate a stronger integration of those functions contributing to the important memory process.

## Methods

### Animals

Wild type male C57BL/6J mice (Jackson Laboratory, stock number: 000664) were group-housed until they were at least 2 months old with unrestricted food and water. They were single-housed after implantation of the microdrive for neural recordings. All experiments were conducted during the animals’ dark phase (i.e., awake phase, 12 h reverse light/dark cycle, lights off at 9 AM). All procedures and animal care were carried out in compliance with guidelines specified by the Institutional Animal Care and Use Committee at the University of Pennsylvania.

### Microdrive assembly

A depth-adjustable microdrive carrying a stimulating electrode and multiple recording electrodes was custom-designed in SolidWorks CAD software to be lightweight and 3D-printed. The stimulating electrode consisted of an insulated tungsten wire that passed through and extended 1.1 mm out of a 28G stainless steel tube (tungsten wire: PFA coated, 51 µm core dia, 102 µm outer dia, A-M Systems). This length was chosen so that when the tip of the tungsten wire reaches the pyramidal cell layer of the dorsal hippocampus, the 28G tube would be in contact with the top part of the cortex directly above the hippocampus. The 28G tube acted as the current return path. Note that in this arrangement, the positive (28G tube) and negative (tungsten wire tip) terminals are spaced apart in the dorsal-ventral direction while remaining concentric with respect to their centers. This arrangement gave a much narrower stimulation artifact compared to twisted bipolar electrodes that had the positive and negative terminals close to each other. To minimize tissue damage, tetrodes were used as recording electrodes. They were made by twisting four insulated Nichrome wires (California Fine wires, Stablohm 800A, material #100189, 12.7 µm core, 20 µm outer dia, heavy polyimide coated). The four terminals were shorted together, and gold plated (to 100 – 300 KΩ impedance), thereby becoming a single recording channel. For easily penetrating the brain, the tips of the stimulating and recording electrodes were cut at a 45 deg angle. Along with the electrodes, ground and reference wires (insulated silver wires, PFA coated, 127 µm core dia, 178 µm outer dia, stripped at the terminals, A-M Systems) were connected to a light-weight electrode interphase board (modified EIB-18, Neuralynx Inc). The electrodes were carried on a shuttle, movable by screws. The microdrive weighed < 1g when assembled (∼1.5g on the mouse head with added dental cement). In several mice, a non-adjustable version of this microdrive was used.

### Surgical procedures

Anesthesia was induced by 3% isoflurane. Once the animals stopped moving, isoflurane was reduced and maintained at ∼2% for another 5 min. Then animals were transferred from the anesthesia induction box to the stereotaxic frame where isoflurane was maintained at 1.75 - 2.25% with progressive reduction of the isoflurane level every hour. The flow rate was 1 L/min. Burr holes were drilled into the skull at AP: −1.7, ML: −1.25 relative to Bregma (Franklin & Paxinos, 2008) for stimulating and at AP: −1.7, ML: +1.25 for the center of the array of the recording electrodes. A ground screw (Antrin Miniature specialists) was threaded on the bone plate over the right cerebellum (interparietal bone) and secured by Loctite superglue. The ground silver wire (insulation stripped for 1.5cm near the tip) was tightly wound around the ground screw and further secured by covering it with silver paint (Silver Print, CRC). A burr hole was made over the left frontal bone and the reference wire (silver ball tip) was inserted under the bone, over the brain surface. When implanting depth-adjustable microdrives, the stimulating and recording electrodes were lowered to DV: −1.0 during surgery. Further adjustments to the final target were made 2-3 days after surgery. For non-adjustable microdrives, the stimulating and recording electrodes were placed at their final locations identified by neural recording during surgery and adjusting the electrodes until the maximal stimulation response on most recording electrodes was obtained. The microdrive was secured to the skull using C & B MetaBond (Parkell) dental cement that bonded the implant to the skull strongly, obviating the need for any additional skull screws.

### Recording procedure

#### Pre-experimental procedures

For mice with adjustable microdrives, after the surgery, we waited for a minimum of 2 days after which the mice were brought to the recording room and the electrodes were lowered by a maximum of 80 µm per day. The characteristic reversal of fEPSP polarity in the pyramidal cell layer along with increased spiking activity was used as a guide to reach the *stratum radiatum*. After that, we waited at least a day to let the electrodes settle. There was one mouse with a non-adjustable microdrive, and that animal was left undisturbed for one week to recover from the surgery and to allow the electrodes to reach a stable location. After this, the mouse was habituated to the recording procedure for at least 3 days before the experiments began. Placing the mouse on top of a tall (6 in) coffee cup (top covered with paper towel), allowed us to quickly connect the head-stage preamplifiers without stress to the mouse. Securing the mouse by holding the microdrive itself, also helped avoid touching the mouse directly. Recordings were done in the dark in the home cage placed inside a sound-attenuating enclosure (Med Associates). The bedding and nest in the home cage were left undisturbed. To avoid tangling of recording cable, the water bottle was removed but a few food pellets were moistened in the cage to minimize dehydration and stress. To stop the mice from climbing out of the home cage, we stacked another cage with its bottom removed, on top of the home cage. For free movement and to reduce the weight added to the head, we used lightweight and flexible recording wires (HS-18 head stage with Cooner wire, Neuralynx) and a motorized commutator (Saturn, Neuralynx) that untwisted the wire. The AC-coupled broad band (0.1 Hz to 16KHz) signal was continuously recorded at a sampling rate of 32 KHz using the Neuralynx Digital Lynx SX system. Stimulus pulses (100µs) were generated by an A-M systems 2100 stimulus isolator with Master-9 pulser (AMPI) periodically giving pulse triggers. Video was acquired at 30 frames/s and synchronized with the neural recording.

#### Neural recording and behavior

At the time of the experiments, mice were ∼ 3 months old (median = 3.3 months, lower-upper quartiles: 2.95 – 3.8 months). For both shocked and control groups, the baseline fEPSP measurement began ∼1.5h (median; IQR: 1.3 – 1.9h) after the house light was turned off. For monitoring synaptic strength, we chose a stimulation current intensity that elicited 40-50% of the maximal response based on an input-output curve, where a set of 5-7 current intensities ranging from 15 to 70 µA were given in a random order, with each current level repeated 3 times. A sigmoidal function was fit to this data from which we computed the 40-50% of maximal response. This current level was used throughout the day. Following this procedure, baseline measurement began using a stimulus pulse (100 µs, monophasic, square shape) that was given every 60s, for 2 hours. After this, the recording was stopped, and the mouse was disconnected. The mouse was then carried in its home cage to the fear conditioning chamber (Coulbourn Instruments), which had been cleaned with 20% ethanol. The conditioning consisted of a 2-minute exploration of the shock chamber after which the mouse received 5 foot-shocks with a 30 s interval between them. Each shock was given at 0.6 mA for 1 s (8-pole scrambled square-wave foot-shock, 40 Hz repetition rate, Coulbourn Precision animal shocker H13-15). After the last shock the mouse remained in the chamber for 30 s. For the no-shock control group, the same above procedure was followed except that no foot shocks were given. To avoid handling-stress to the mouse, after the shocking procedure, instead of capturing the mouse by hand, we used a cardboard scoop (“shepherd shacks” used for mice) on to which the mice easily entered. We returned the mouse to his home cage for recording and continued the stimulation pulses as in the baseline for another 3 hours. For the shocked group, memory retention was tested either 24h later (n = 12) or after 7 (n = 9) or more days (n = 3, one mouse each after 8, 10 and 27 days) by measuring freezing (FreezeFrame, Actimetrics) in the original shocked chamber for 5 min. The mouse group that was tested 7 or more days after conditioning is referred to as the 7-day post-conditioning group in the Results section. Although not blind to the experimental conditions, the same experimenter performed all the microdrive implantations and handled all mice the same way during the experiments.

### Histology

To localize the electrode tips, we first anesthetized the mouse with ketamine-xylazine mixture and passed 30 µA positive current for 8-10 s through the electrode tips with the ground or reference as the current return path. Cardiac perfusion was then performed with 10% formalin (in phosphate-buffered saline) fixative. The mouse head with the microdrive implant was submerged inside the fixative at 4 C for at least 1 day. After that, leaving the region of skull cemented to the implant intact, the rest of the tissue surrounding the brain was carefully removed. Finally, the brain was gently separated out of the skull/implant and fixed in 10% formalin for another day. The brain was then sliced coronally at 50 µm thickness in a vibratome.

### Data analysis

All data analyses were carried out using custom-written MATLAB code and the large-scale data management framework – DataJoint (Yatsenko et al., 2015). Statistical testes were done using MATLAB’s Statistics Toolbox unless otherwise noted. The experimental quantities (e.g. length of sleep episodes) in the Results section are reported either as mean ± 1 Standard Error (SE) or as median with Inter Quartile Range (IQR). When a quantity (e.g. NREM state length) did not follow normal distribution as determined by the Anderson-Darling test, we opted to use the median. Further, for simplicity, when only one of two related quantities (e.g. NREM state length and REM state length) did not follow normal distribution, we report median for both. Sample size (n) refers to number of mice unless otherwise stated.

#### Slope measurement of the fEPSP

First, the broadband signal was down-sampled to 16 KHz and filtered at 0.1 – 5000 Hz using MATLAB’s decimate routine. The fEPSP stimulation response traces were then extracted using the stimulation onset timestamps. To compute the slope of the initial downward deflection, a second derivative of the trace was computed. This derivative had a single prominent peak at the time when the downward deflection was roughly halfway down. We then placed a 500 ms window centered on the time of the derivative’s peak and computed the slope from the raw trace in this time window.

#### Motion estimation

The mouse’s position was detected online by thresholding the red LED tracker lights attached to the head-stage preamplifiers. A motion index (*m*) was computed using the following heuristic formula based on the robust estimate of standard deviation (Quiroga et al., 2004) of the x and y coordinates (***x***, ***y***) of 5 s contiguous time windows (5s x 30fps = 150 pairs of data points):

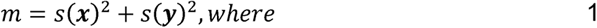

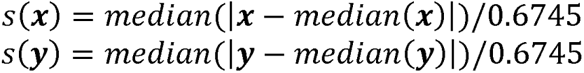

The median-based robust standard deviation was employed for the following reasons. The position tracker software sometimes mistakenly detected the reflection of the tracker LED on the sidewalls of the mouse cage as the position of the mouse. At other times, the recording cable or the nest of the mouse blocked the LED from the view of the camera resulting in loss of mouse position; the tracking software assigns a position of (0,0) in these cases. All these artifacts result in sudden jumps in the otherwise smoothly changing values of the x and y coordinates. By employing median, we minimized the contribution of these artifacts thereby avoiding overestimation of the motion index.

#### Identification of behavioral states

Behavioral states were identified by combining mouse motion information with the hippocampal theta/delta band power ratio (TDR). Power in the theta band (4-11Hz) and delta band (1-4 Hz) was computed in 5 s contiguous time windows using *bandpower* routine of MATLAB (MathWorks, MA). TDR was averaged across electrodes within each mouse. The motion index (MI) was also computed in 5 s contiguous time windows as described in the previous section. We detected REM and NREM sleep states by thresholding MI and TDR as follows. For threshold values, we first computed the standard deviation (SD) (as in eq 1) of MI and TDR for each pre- and post-conditioning recording session. Time segments with motion index (smoothed with a Gaussian kernel with SD = 5s) values below 0.01 MI-SD were treated as putative sleep segments. We used TDR to identify REM and NREM periods within these tentative sleep segments. Although TDR is characteristically lower during NREM than during REM and awake state, we found that, for the purpose of NREM state identification, the TDR variance proved much easier as it reduces considerably during NREM (Figure 3C and Supplemental Figure S2C). To utilize this property for sleep state identification, we computed a moving variance (MV) analogous to a moving average by computing SD (as in eq 1) of TDR values in 60s moving windows and then squaring the SD. Putative sleep segments with MV less than 0.5 TDR-SD were classified as putative NREM periods whereas those with MV more than 1 TDR-SD were classified as putative REM periods.

The thresholding sometimes spuriously identified short isolated time segments as NREM or REM. To exclude these, we set minimum segment lengths of 15 and 30s for REM and NREM respectively. Rarely, spurious REM segments were detected that were not closely preceded by a NREM period. Since in wild type mice NREM precedes REM (Chemelli et al., 1999), we excluded such spurious REM segments by requiring NREM to precede REM within 180s. Rarely, we excluded short (< 30s) isolated putative NREM segments without nearby (<5 min) NREM or REM segments. Time segments with MI values crossing the MI threshold for less than 5s were joined with the adjacent sleep segments. Interruptions of sleep by brief arousal/awakenings (∼5-60s) occur naturally in mice (Chemelli et al., 1999; Franken et al., 1999; Huang et al., 2006; Léna et al., 2004; Lima et al., 2017; Perez-Atencio et al., 2018; Watson et al., 2016). These periods were labeled as ‘brief awakenings’ and were not included in the ‘awake’ state when comparing the effect of NREM on fEPSP with that of awake state (Figure 3E and Figure 4C-E). The NREM-mediated enhancement of fEPSP slope may extend into the brief awakening periods. Hence, when measuring learning effects, as a conservative measure, we combined REM, NREM and the brief awakenings that occur within 90s of each other into a single ‘sleep episode (Supplemental Figures S2E, S3 and S5). In addition, the transition period from awake to NREM state and vice versa, may have a partial enhancing effect on fEPSP as does NREM state. To minimize this effect confounding the learning effect, we extended the boundaries of the sleep episode to include the transition period. We defined the transition time point as the time when there was a significant change in the mean (*findchangepts* routine of MATLAB) of the TDR values in a time window (< 4 min) centered at either boundary of the sleep episode. The fEPSP data points corresponding to the sleep episode were removed when measuring learning effects (Figure 6, Figure 7D and Supplemental Figure S1C). Time periods outside the sleep episodes were treated as the ‘awake state’. However, when comparing the effect of fear conditioning on sleep itself (Figure 4B), only REM and NREM segments were included, i.e., brief awakenings and transition periods were excluded.

The above sleep detection parameters were determined heuristically by how well they classified sleep states. Once determined, the same values were used for all recording sessions of all experimental conditions. Sleep state identification was carried out by custom written MATLAB code automatically without any further manual intervention.

For the spectrogram in Figure 3B, we computed a continuous wavelet transform in MATLAB using complex Morlet wavelets (see the Supplemental materials).

#### Colormaps

Two-color (red and blue) divergent colormaps were created using the methods described in the literature (Moreland, 2009). This colormap ensures that the order of data values is reflected in the order of perceived colors and that the rate of color change is in proportion to the rate of change in data value. We allocated the number shades of red and blue in proportion to the range of data values above and below the neutral data value (e.g.,100 for normalized fEPSP slopes). This color code ensured that the neutral data values were depicted in the neutral color (gray) of the colormap.

#### Estimation of NREM/awake ratio

To examine if the NREM mediated enhancement of synaptic strength outlasts the sleep episode (Figure 4E), for each mouse, we computed the NREM-to-awake fEPSP slope ratio separately for each of the sleep-awake segment pairs, and then we averaged these ratios: median length of sleep episode = 12.8 min (IQR: 11 – 15.1 min); median length of awake state = 26.4 min (IQR: 19 – 36 min), n = 17 mice. When the awake state was restricted to a maximum of 15 min post-sleep, the median length of the selected awake states was 15 min (IQR: 11.8 – 15 min). Only mice with at least one sleep-awake pair with at least 5 fEPSP measurements in each state were included in the analysis.

#### Estimation of sleep-mediated renormalization of synaptic strength

In the post-conditioning sessions, we first selected sleep episodes that had at least 5 min of NREM state. Then, as opposed to selecting the complete awake period that precedes or follows a sleep episode, we selected two narrow 5-min awake-state time windows - one that immediately precedes (pre-sleep) and another that immediately follows (post-sleep) a given sleep episode (Figure 4F, Top). These shorter 5-min windows were chosen to favorably detect any sleep-induced reduction in synaptic strength. Because synaptic strength ramps up during the awake state (Vyazovskiy et al., 2008), the synaptic strength should reach its maximal value in the last part (our 5 min window) of the awake period that precedes the sleep episode. On the other hand, since synaptic strength ramps down during the sleep state, in the awake state that immediately follows, the lowest synaptic strength will be detected at the initial part of the awake state. Hence the 5-min windows we selected should favorably capture any renormalization by sleep. This time window is also comparable to the 1-3 min window used previously (Vyazovskiy et al., 2008). We did not allow the awake period windows to overlap. When the post-sleep awake state window of one sleep episode overlapped with the pre-sleep awake state window of the next sleep episode, the sleep episode with the shorter NREM period was excluded. We computed the average fEPSP slope within the post- and pre-sleep awake state windows and obtained the post/pre ratio of fEPSP slope. The ratios obtained from multiple sleep episodes (1-3 per mouse) within a mouse were averaged. The part of pre- and post-sleep awake state used for analysis was fixed at a length of 5 min, whereas the median length of the full pre-sleep awake state was 36 min (IQR: 16 – 40.8 min) and that of post-sleep awake state was 24 min (IQR: 17.2 – 30.3 min), n = 15 mice.

#### Estimation of instantaneous speed

Using the x- and y-coordinates from the LED-based position tracking, we computed the instantaneous speed of the mouse at each time point as described previously (Hen et al., 2004). Briefly, the instantaneous speed was computed as a first derivative of a second-degree polynomial fit separately to the x- and y-coordinates that had been smoothed by the LOWESS (Locally Weighted Scatter plot Smoothing) method. In the LOWESS algorithm, we iteratively fitted a second-degree polynomial to the data points of a moving window of 15 video frames (equivalent to 0.5 s). The polynomial fitting was done using a weighted linear regression where smaller weights were given to points away from the current time point and to points that were farther from the fitted curve (i.e. bigger residuals). Animal speed was then computed as the square root of the sum of the squares of the x- and y-component instantaneous speeds. These steps were performed using custom-written MATLAB code and using built-in routines *lscov, polyder* and *polyval*.

#### Correlation between instantaneous speed and fEPSP slope

Repeated measures correlation between instantaneous speed and fEPSP sleep was computed using the “rmcorr” package (Bakdash & Marusich, 2017) in the statistical software R.

#### Inclusion criteria for mice and electrodes

Electrodes that were outside the CA1 region were excluded. In addition, to get reliable slope estimates, electrodes in the CA1 that had weak stimulation responses (fEPSP peak magnitude less than ∼0.5 mV) were also excluded. Similarly, electrodes that landed near the pyramidal cell layer were also excluded because they occasionally showed a popspike, which makes the slope measurement unreliable. In addition, electrodes with slope values that showed systematic drift (i.e., were not stable) during the baseline (see the Baseline stability testing section below) were excluded (0/62 in the cFC group and 2/25 in the sham-conditioning group). Mice without any stable electrodes were also excluded (0/24 in the cFC group and 1/13 in the sham-conditioning group).

#### Baseline stability testing

A stable baseline of synaptic strength is a prerequisite for a valid interpretation of the post-conditioning synaptic changes. Therefore, we defined the baseline stability at the individual electrode and at the population level. First, at the individual electrode level, if the baseline drifts up in some electrodes and drifts down in others, they can cancel out each other at the population level, resulting in a ‘steady’ average baseline. Hence, we tested stability of individual electrodes as described in the next section. It is also possible that in most or all electrodes there may be a small drift in the same direction. In that case, the electrode-level test will fail to identify drift adequately, whereas at the population level, these small effects will add up to be easily detectable. Therefore, we also performed stability tests on the average across electrodes (see *Statistical tests for learning effect* section).

#### Single electrode level stability test

To test for baseline stability in a principled way, we used the following statistical procedure similar to the Kolmogorov-Smirnov (KS) statistic that quantifies the distance between the empirical distribution function of a sample and the cumulative distribution function of a reference (null) distribution. The normalized fEPSP slope data (Supplemental Figure S6A, sleep segments removed) ordered in time is the sample distribution, the cumulative sum of which forms the empirical distribution function (*F*, Supplemental Figure S6C, black). Then, we created a reference distribution that denotes a perfectly stable baseline with no change in fEPSP slope over time. As an ideal stable baseline would have 100% as the value of normalized fEPSP slope at each time point, we created such a sequence where each element was 100 (Supplemental Figure S6B). The length of this sequence was the same as that of the experimentally measured fEPSP slope data. The cumulative sum of the ideal stable baseline data points formed the reference distribution function (*F_n_*, Supplemental Figure S6C, gray). As a test statistic (*D*), similar to the KS statistic, we computed the maximal absolute distance between the two distribution functions (*F* and *F_n_* Supplemental Figure S6C). To find if this statistic is significant, we created a null distribution by bootstrapping, where we repeated the computation of the test statistic (*D*^*^, Supplemental Figure S6E) 10,000 times, each time randomly picking the same number of fEPSP slope data points (Supplemental Figure S6D) as the original data (resampling with replacement). Such shuffling results in a sequence of fEPSP slope data that does not have systematic drift over time but preserves the noise in the data (Supplemental Figure S6D). We then computed the fraction of bootstrapped D* values (Supplemental Figure S6F) that were above D, as the p-value. We treated any p-value less than 0.05 as indicative of a significant drift of the slope data. Electrodes with such fEPSP slopes were considered unstable.

#### Statistical tests for learning effect

We sought out a statistical procedure that can test the stability of the baseline and the learning-related changes more comprehensively with the ability to handle missing data. We opted for the linear mixed model (LMM) formulated for analyzing interrupted time series. We treated fEPSP slope data points as an interrupted time series (Wagner et al., 2002) where the baseline time series is “interrupted” by fear conditioning. A linear fit was applied to the baseline and post-conditioning fEPSP slopes (segmented linear regression, see Supplemental materials). The linear fit contained two types of parameters – intercept and gain. Note that, the term ‘gain’ is used instead of ‘slope’ to refer to the slope of the line fit. This naming avoids the slope of the fEPSP being confused with the slope of the line fit.

The intercept denotes the level on the y-axis at which the fEPSP slopes start at the beginning of the baseline or at the beginning of the post-conditioning period. Learning may result in an abrupt “jump” or increase in the fEPSP slopes at the beginning of the post-conditioning period (Figure 2F, Figure 6A and Supplemental Figure S1B-C). A significant increase in the intercept parameter in the post-conditioning period relative to baseline thus reflects a significant increase in post-conditioning fEPSP slopes.

The gain value indicates whether the fEPSP slopes remain stable (zero gain), grow (positive gain) or decline (negative gain) over time. For example, for the baseline period, a gain value that is insignificantly different from zero indicates that the baseline is stable (Figure 2F, Figure 6, Figure 7D and Supplemental Figure S1B-C). A positive gain value of the fit to the post-conditioning period will indicate that the fEPSP slopes grow further with time relative to those of the baseline.

LMM analyses were done in the SPSS statistical software package PASW-18. Briefly, the mice were treated as random effects, and time points of baseline and post-conditioning periods were treated as fixed effects. The normalized fEPSP slope was used as the dependent variable. Although for the display purposes the raw data points were 5-min binned in Figure 2E-F, Figure 6, Figure 7B & D and Supplemental Figure S1, for statistical tests, the non-binned raw normalized fEPSP slopes were used. For example, for statistical tests of Figure 2F, the baseline had 120 data points per mouse. Note that we treated the first 10 min of the post-conditioning period as a transition period. To avoid modeling this transition that would require additional model parameters, we excluded the data (10 data points) corresponding to this period during statistical tests.

### Data availability

Raw data and MATLAB code associated with this study are available at https://doi.org/10.5061/dryad.sj3tx965f.

## Acknowledgements

We thank Shannon Wolfman and Daniel Kalamarides for their thorough feedback on the manuscript, Dimitri Yatsenko for help with statistical methods, Marion Scott for technical and administrative assistance, Tiffany Brown-Mangum and Tyisha Hundley for their help with animal husbandry. This work was supported by National Institutes of Health grants R01AA026267 (National Institute on Alcohol Abuse and Alcoholism, J.A.D.) and R01NS021229 (National Institute of Neurological Disorders and Stroke, J.A.D.). This work also was supported by a generous award from the Chernowitz Medical Research Foundation.

## Supplemental materials

### Computing spectrogram based on Morlet wavelet transform

For the spectrogram in Figure 3B, we computed a continuous wavelet transform in MATLAB using complex Morlet wavelets (*ψ*(*t*)) defined by the following equation:

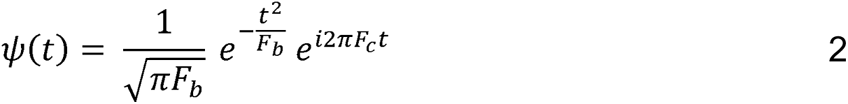

In this equation, *t* is time, *F_c_* is the center frequency of the wavelet, and *F_b_* controls how fast the wavelet decays in the time domain. The energy spread of the wavelet in the frequency domain is a Gaussian and the parameter *F_b_* is inversely related to the variance 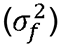 of this Gaussian by the following relationship:

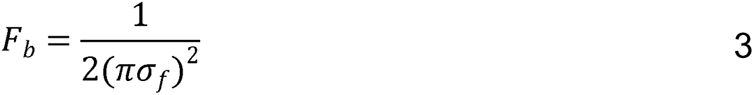

To compute the spectrogram in Figure 3B, we created a series of 30 wavelets (half width of 2.5 s) by varying *σ_f_* (0.125 to 2.5Hz) and *F_c_* (1 to 30Hz) both on a logarithmic (10 based) scale. Each wavelet was then normalized to have unit norm. The raw data (sampled at 32 KHz) was first down-sampled to 320 Hz, and its wavelet transform was then computed in 30s-long contiguous data segments with a 2s buffer at either end of each segment. The resulting spectrogram was z-scored across the frequencies at each time point.

### Linear mixed model analysis (LMM)

We describe the LMM using an example with 120 fEPSP slope measurements during baseline and 180 in post-learning period. The subjects (mice) are treated as random effects; instantaneous speed of the animal and the time points at which the speed and fEPSP slopes were measured at the different segments (see below), are treated as fixed effects. The fEPSP slope (S*_tj_*) of an electrode from a mouse (*J*) at a given time point (*i*) is modeled as follows:

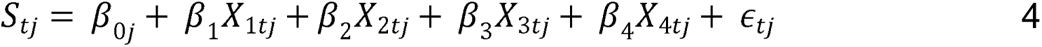

Where,*β*_0_ denotes subject-specific intercept and *ϵ*_tj_ denotes error associated with slope *S_tj_*. Time (in minutes) since the beginning of the experiment is denoted by *X*_1*tj*_ where

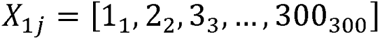

The subscripts on the right-hand side of the above equality denote time *t*. At each time point, the pre- and post-learning conditions are coded as 0’s and 1’s respectively (dummy coding) by *X*_2*tj*_ where

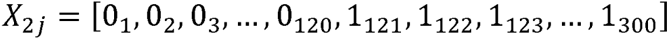

Time since “interruption” (fear conditioning) is denoted by *X*_3*tj*_ where

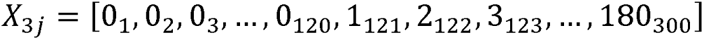

The animal’s instantaneous speed (transformed by 2-based log), from the beginning of the experiment is denoted by *X*_4*tj*_ where

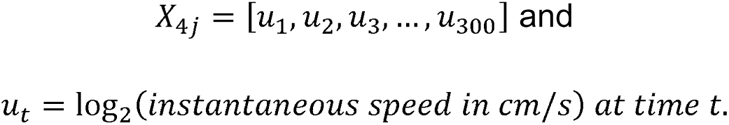

*β*_0_, *β*_1,_ *β*_4_ can be interpreted by applying the values of *X*_1_, *X*_2_, *X*_3_ and *X*_4_ in Eq.4 For the baseline period(t ≤ 120), we obtain

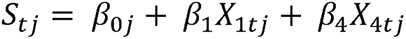

For both baseline and post-learning period (see below), a significant *β*_4_ parameter would indicate that the speed of the mouse contributes significantly to the measured fEPSP slope. Hence a non-significant *β*_1_ parameter would indicate that there is no significant drift in the baseline after accounting for any speed effect. Similarly, for the post-learning period (t > 120)

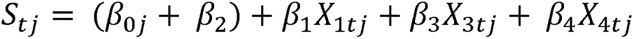

Hence, a significant *β*_2_ parameter would indicate that there is significant shift in the level of post-conditioning data points relative to baseline after accounting for any speed effect. A non-significant *β*_3_ would suggest that the post-learning trend is not significantly different from that of baseline after accounting for any speed effect. That is, the interruption did not significantly alter the ongoing trend (if any) in the baseline. Note that for Figure 2F, the speed term *β*_4_ *X*_4_ was not included in the model.

For directly comparing the learning effect in the fear-conditioned group against that of the sham-conditioned group, the model complexity was reduced by first removing components containing gain parameters - *β*_1_ *X*_1_ and *β*_3_ *X*_3_ (see Eq. 4) and adding a component *β*_5_ *X*_5_ where *X*_5_ codes data points from shocked and control (sham) groups as 1’s and 0’s respectively. In addition, a term for interaction between pre-post and shocked-sham conditions - *β*_1_ (*X*_2*tj*_*X*_5*tj*_ was included.

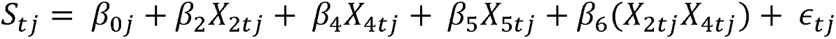

A significant *β*_6_ parameter would indicate that the change in the fEPSP slopes in the post-conditioning period relative to baseline, depends on the shocked-sham condition, after accounting for any speed effect.

## Figure legends for Supplemental Figures

**Figure S1.**
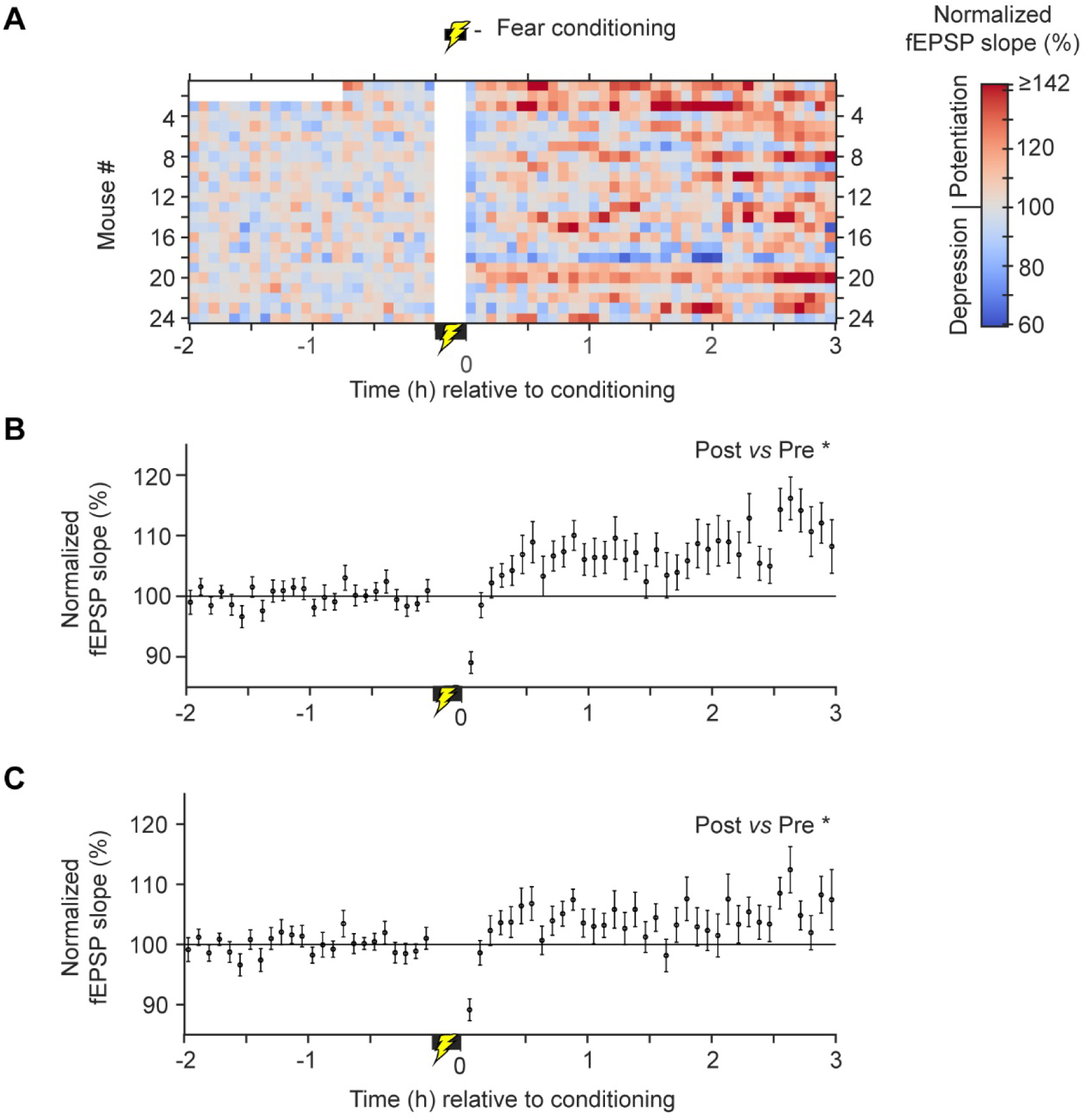
In vivo synaptic strength modifications associated with fear conditioning measured by averaging electrodes within each mouse. ***A,*** Heat map of 5-min binned slopes of fEPSP responses that were first normalized to baseline and then averaged across electrodes within each mouse. Each row shows data from a mouse (n = 24). The conditioning period is marked by the yellow lightning bolt symbol near time zero. Mice #1 & 2 had only 45 min of baseline data. Mice # reflects ascending chronological order of recording dates. ***B,*** Across-mouse average of 5-min binned normalized slopes. ***C,*** As in ***B*** except that fEPSP slopes corresponding to sleep segments were removed before averaging. Error bars are SEM. * p < 0.05, significant increase in fEPSP slopes in post-conditioning period relative to baseline.

**Figure S2.**
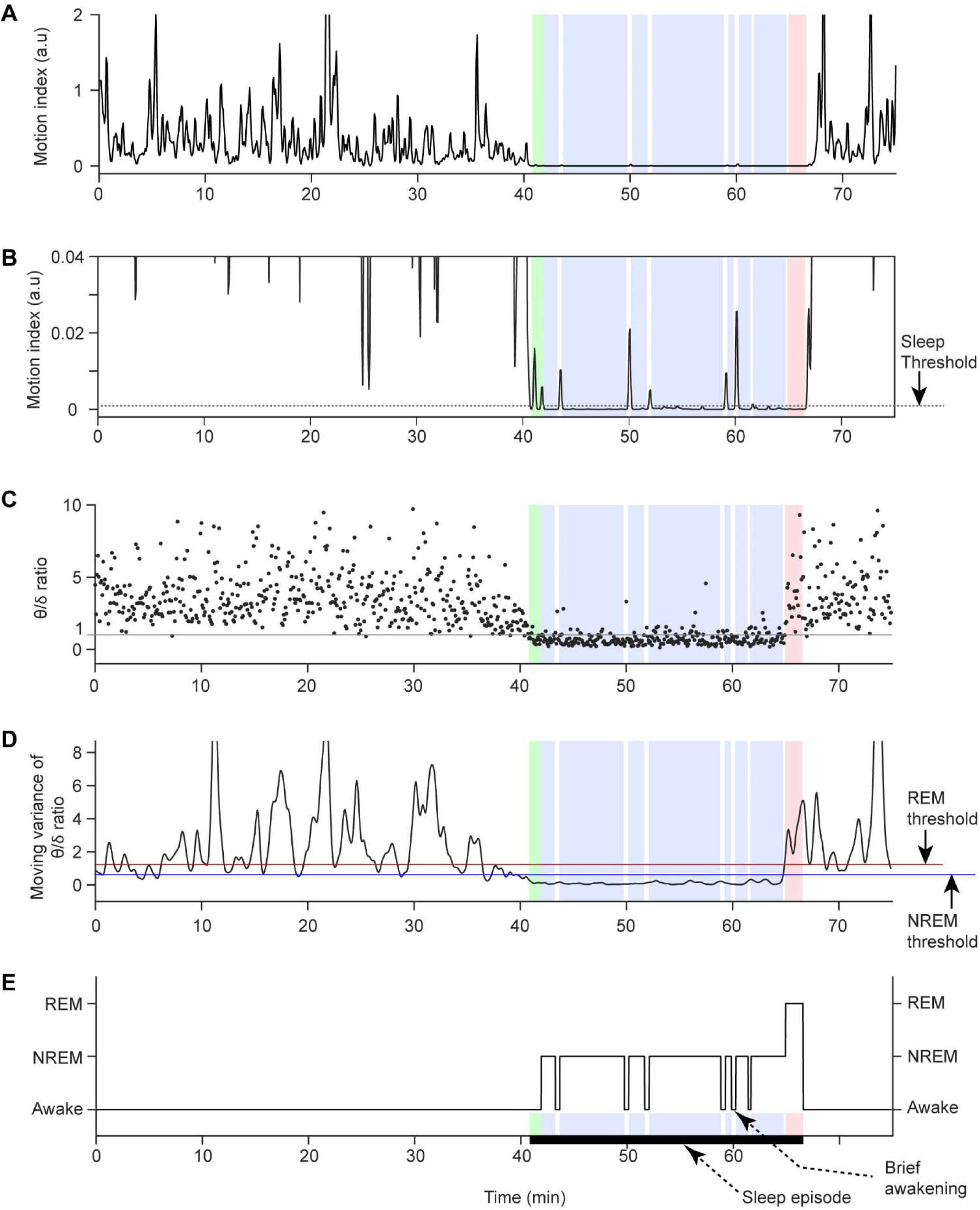
Sleep state identification procedure. ***A,*** Motion index (MI, smoothed with a Gaussian kernel) of the first 75 min of post-conditioning period of an example mouse. The colored regions show different sleep segments (see below). ***B,*** MI data in ***A*** enlarged to show sleep detection threshold (dotted gray line). The threshold is 0.01 times the standard deviation (SD) of the MI. Time segments with MI values below the threshold were putative sleep segments. Note the brief awakenings marked by small animal movements in the time period between minute 40 and 70. Outside of the colored sleep region, the movement is mostly off this movement scale. ***C,*** Theta/delta power ratio (TDR). Note the marked decrease in the TDR values and their variance during the putative sleep period. ***D,*** Variance of TDR computed in a moving window (60s width) and then smoothed with a Gaussian kernel. REM sleep threshold (red line) is 0.5 SD of TDR and NREM sleep threshold (blue line) is 1 SD of TDR. ***E,*** Hypnogram. Putative sleep segments (from B) with moving TDR variance below the NREM threshold were identified as NREM sleep segments (blue shaded boxes). Putative sleep segments with moving TDR variance above the REM threshold were treated as REM sleep segments (red shaded box). The green shaded box indicates part of the period of transition from awake to NREM state. The periods corresponding to the transition, NREM, REM and the brief awakenings that occur in between the segments of NREM and REM, were grouped into a single sleep episode (thick black line over the x-axis).

**Figure S3.**
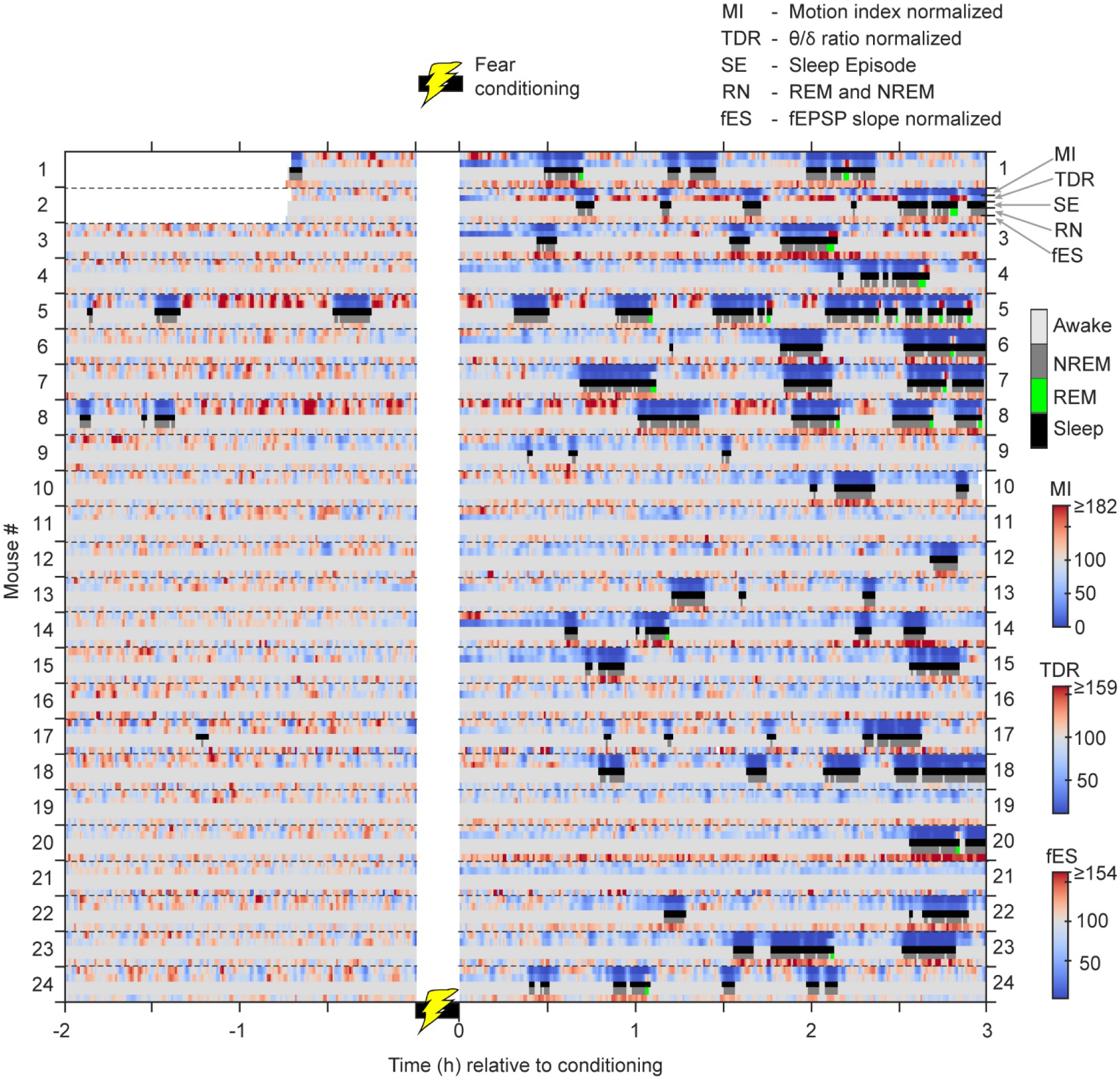
Normalized fEPSP slopes (fES) of the most potentiated electrode set of the fear conditioned mice, aligned with sleep episodes (SE, black). Motion index (MI, square root transformed) and theta/delta band power ratio (TDR) were normalized to their respective baseline for each mouse. When multiple electrodes were present in a given mouse, normalized TDR values were averaged across electrodes. NREM (dark gray) segments were identified by sustained decrease (cold blue) in MI and TDR. REM (green) segments were identified by sustained decrease in MI and increase (warm red) in TDR. Multiple NREM and REM segments (RN) separated by less than 1.5 min were grouped into a single sleep episode (black). MI and TDR were computed in 5s contiguous time windows and smoothed with a Gaussian kernel (30s standard deviation). fEPSP slopes (1 pulse/min) were not binned but smoothed with a Gaussian kernel (45s standard deviation). Dotted horizontal lines mark boundary of data set of each mouse.

**Figure S4.**
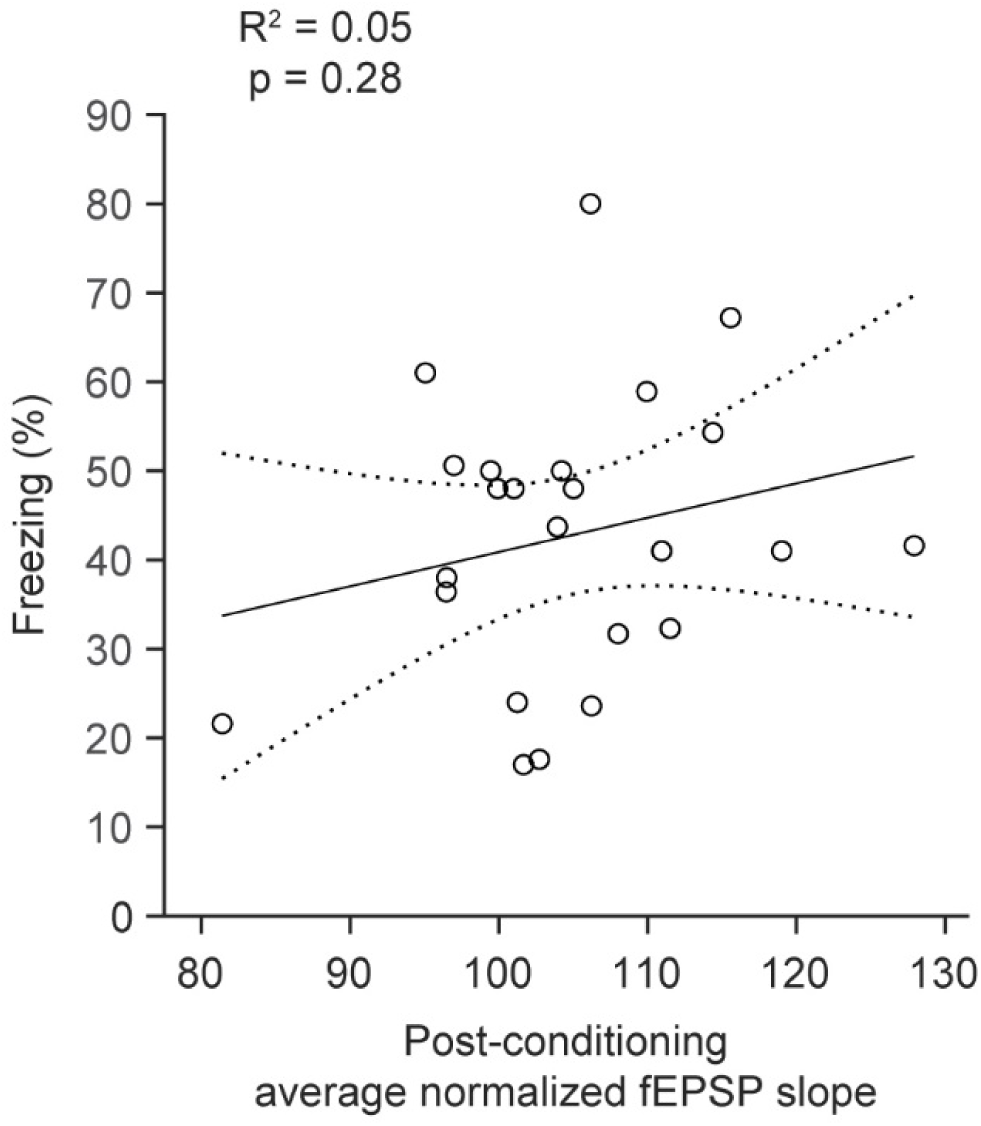
Correlation between behavioral measure of memory (percent freezing) and physiological measure of plasticity (normalized fEPSP slopes averaged over the post-conditioning period). Each data point represents a mouse (n = 24). Linear model fit (black line) is shown along with the 95% confidence bounds (dotted traces).

**Figure S5.**
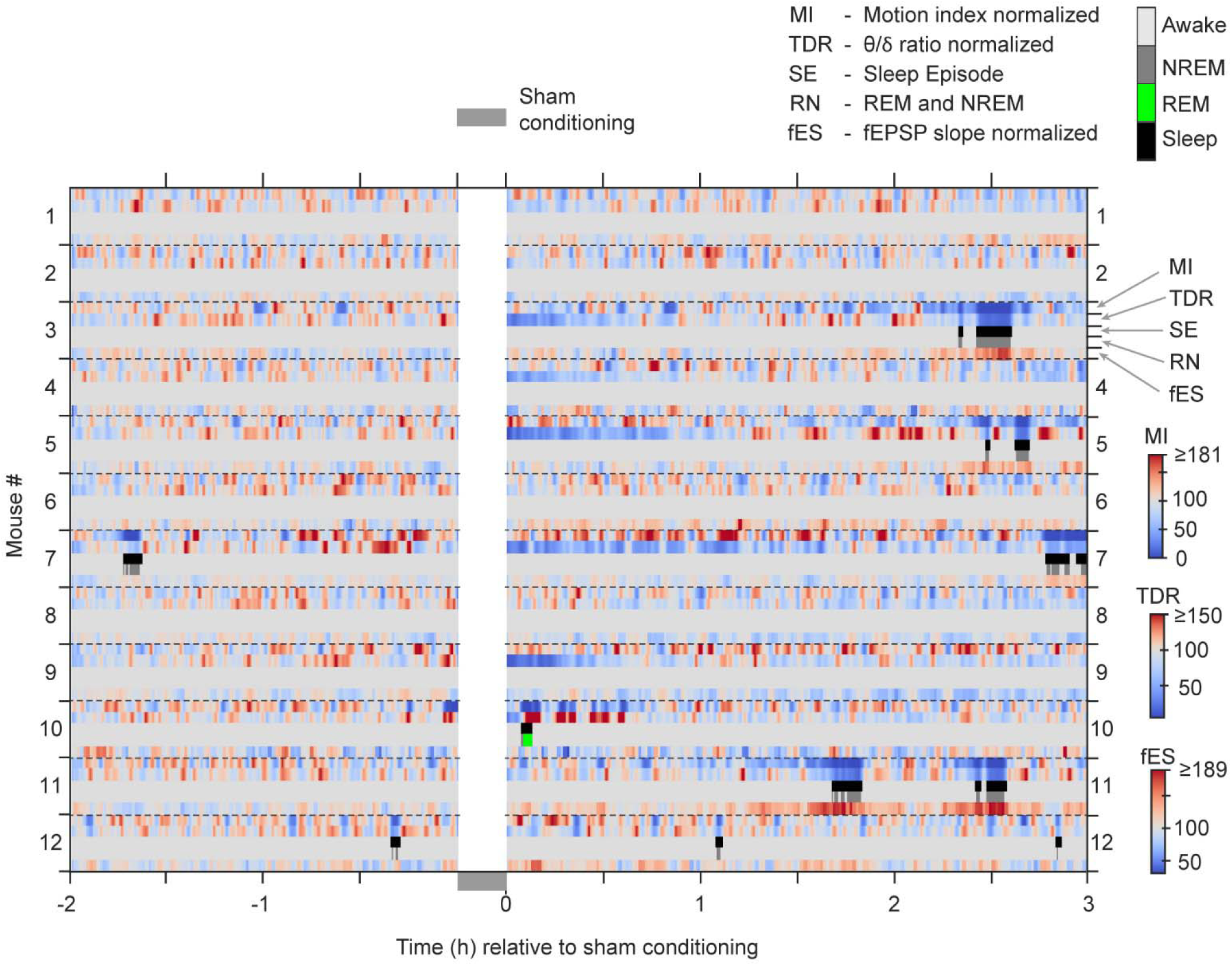
Normalized fEPSP slopes (fES) of the most potentiated electrode set of the sham conditioned mice, aligned with sleep episodes (SE, black). Motion index (MI, square root transformed) and theta/delta band power ratio (TDR) were normalized to their respective baseline period values for each mouse. When multiple electrodes were present in a given mouse, normalized TDR values were averaged across electrodes. NREM (dark gray) segments were identified by sustained decrease (cold blue) in MI and TDR. REM (green) segments were identified by sustained decrease in MI and increase (warm red) in TDR. Multiple NREM and REM segments (RN) separated by less than 1.5 min were grouped into a single sleep episode (black). MI and TDR were computed in 5s contiguous time windows and smoothed with a Gaussian kernel (30s standard deviation). fEPSP slopes (1 pulse/min) were not binned but smoothed with a Gaussian kernel (45s standard deviation). Dotted horizontal lines mark the boundary of the data set of each mouse.

**Figure S6.**
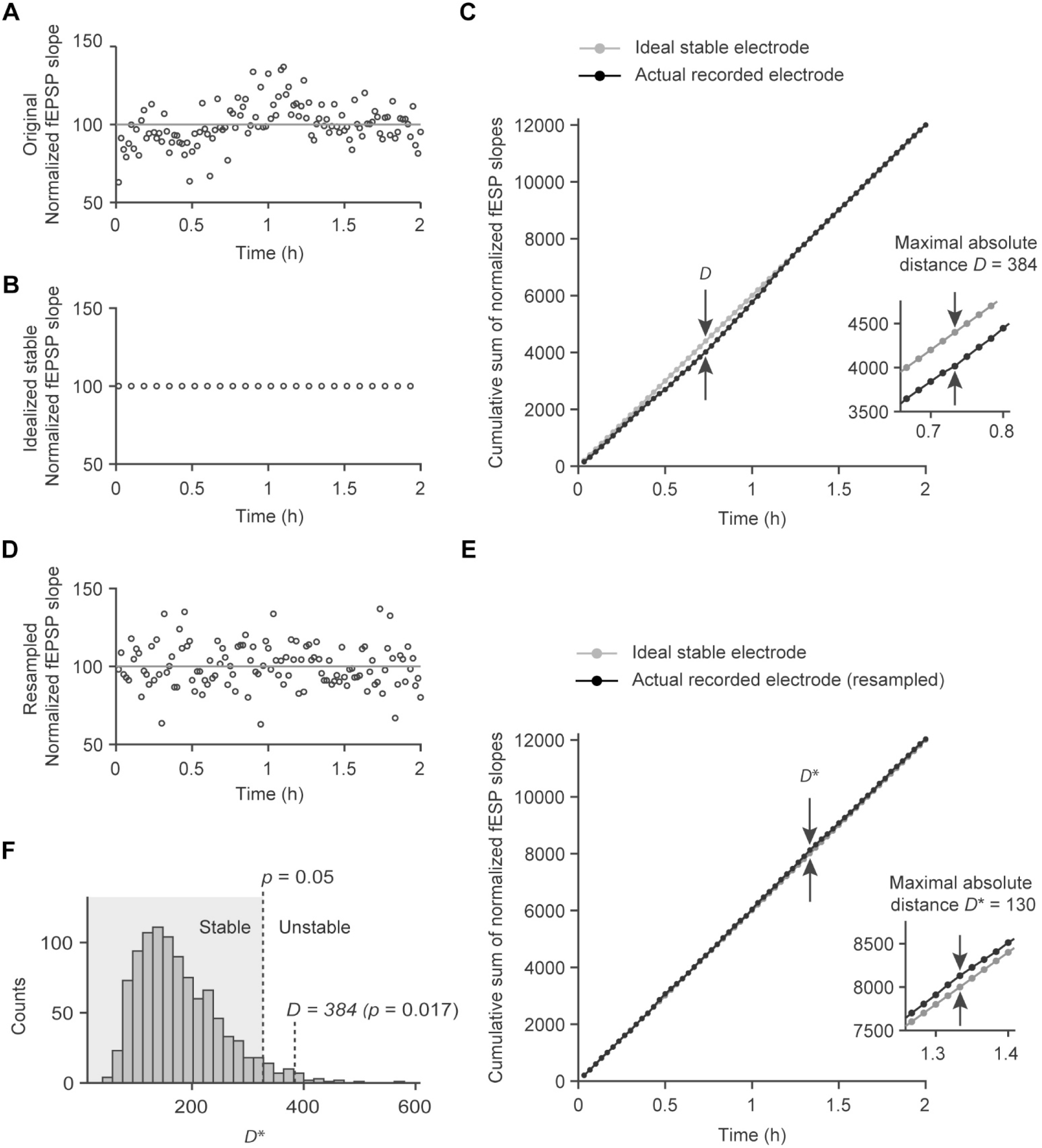
Statistical method for testing electrode stability. ***A,*** Baseline fEPSP slopes (normalized to mean) recorded from a likely unstable electrode. ***B,*** Hypothetical ideal stable baseline where each data point is set to 100. Note that only a subset of data points is shown although for analysis the number of data points was matched to that in ***A. C,*** Cumulative sum of the fEPSP slopes from panel ***A*** (black) and panel ***B*** (gray). For clarity, only every other data point is shown. The largest vertical separation between the two cumulative sums is indicated by the opposing vertical arrows. The magnitude of this separation is defined as the maximal absolute distance, *D*, as shown enlarged in the inset. To test if separation as big as *D* could have come from a stable electrode, a null distribution representing a stable electrode was created by resampling the original fEPSP slopes (bootstrapping). ***D,*** One instance of resampling (with replacement) of data points of panel ***A***. Note that any trend in the data is now lost due to the resampling. ***E,*** Cumulative sums of one instance of resampled fEPSP slopes (black) and of idealized stable fEPSP slopes (gray). The largest vertical separation between the two cumulative sums (*D*\*) represents how large *D*\* can go simply due to noise in the data even though the electrode is stable. ***F***, Null distribution histogram of *D*\* values obtained by 1000 iterations of resampling the original fEPSP slopes (as in ***D***) and then computing *D*\* (as in ***E***). Under this null distribution of a stable electrode, a *D* value as large as 384 (computed in ***C***) is unlikely (p = 0.017). With a threshold of p = 0.05 for deciding stability, the electrode in ***A*** is deemed unstable.

